# CSPP1 stabilizes growing microtubule ends and damaged lattices from the luminal side

**DOI:** 10.1101/2022.06.23.497320

**Authors:** Cyntha M. van den Berg, Vladimir A. Volkov, Sebastian Schnorrenberg, Ziqiang Huang, Kelly E. Stecker, Ilya Grigoriev, Sebastian Patzke, Timo Zimmermann, Marileen Dogterom, Anna Akhmanova

## Abstract

Microtubules are dynamic cytoskeletal polymers, and their organization and stability are tightly regulated by numerous cellular factors. While regulatory proteins controlling formation of interphase microtubule arrays and mitotic spindles have been extensively studied, the biochemical mechanisms responsible for generating stable microtubule cores of centrioles and cilia are poorly understood. Here, we used in vitro reconstitution assays to investigate microtubule-stabilizing properties of CSPP1, a centrosome and cilia-associated protein mutated in the neurodevelopmental ciliopathy Joubert syndrome. We found that CSPP1 preferentially binds to polymerizing microtubule ends that grow slowly or undergo growth perturbations and, in this way, resembles microtubule-stabilizing compounds such as taxanes. Fluorescence microscopy and cryo-electron tomography showed that CSPP1 is deposited in the microtubule lumen and inhibits microtubule growth and shortening through two separate domains. CSPP1 also specifically recognizes and stabilizes damaged microtubule lattices. These data help to explain how CSPP1 regulates elongation and stability of ciliary axonemes and other microtubule-based structures.

## Introduction

Microtubules are dynamic cytoskeletal polymers that serve as tracks for intracellular transport and drive chromosome separation during cell division. The majority of cellular microtubules turn over rapidly because microtubules frequently switch between phases of growth and shortening (Desai and Mitchison, 1997). Proteins controlling microtubule dynamics in interphase and mitosis have been studied in great detail by a combination of genetic, cell-biological, biochemical and biophysical experiments (Akhmanova and Steinmetz, 2015; Gudimchuk and McIntosh, 2021). In particular, in vitro reconstitution studies with purified components have been instrumental for understanding the mechanisms underlying the activity of these proteins (Bieling et al., 2007; Gell et al., 2010). However, cells also form stable microtubule-based structures, such as centrioles and cilia. Multiple molecular players responsible for biogenesis of centrioles and cilia have been identified through genetic and cell-biological approaches, but their biochemical properties and their effects of microtubule dynamics are very poorly understood, because most of them have never been investigated using purified proteins.

Here, we focused on the centrosome/spindle pole associated protein 1 (CSPP1) (Patzke et al., 2005; Patzke et al., 2006). Previous work established that CSPP1 binds to spindle poles and central spindle during mitosis and to ciliary axonemes, centrosomes and centriolar satellites in interphase (Asiedu et al., 2009; Frikstad et al., 2019; Patzke et al., 2005; Patzke et al., 2010; Patzke et al., 2006). CSPP1 accumulates at ciliary tips, interacts with several other ciliary tip proteins, contributes to ciliogenesis and controls axoneme length. Loss of CSPP1 leads to the formation of shortened cilia and impaired Hedgehog signaling, which depends on ciliary function (Frikstad et al., 2019; Latour et al., 2020; Patzke et al., 2010). Mutations in genes encoding CSPP1 and its ciliary binding partners lead to defects in ciliogenesis and result in a range of ciliopathies, such as the neurodevelopmental disorder known as Joubert syndrome, or the more severe Meckel-Gruber syndrome with multiple developmental abnormalities (Akizu et al., 2014; Latour et al., 2020; Shaheen et al., 2014; Tuz et al., 2014).

While the tissue and cellular phenotypes associated with CSPP1 defects have been analyzed in some detail, very little is known about its mechanism of action. To close this knowledge gap, we have performed in vitro reconstitution experiments to investigate the impact of full-length CSPP1 and its individual domains on microtubule dynamics. We found that CSPP1 specifically associates with growing microtubule ends when they undergo a growth perturbation and enter a pre-catastrophe state.

CSPP1 stabilizes such ends and induces microtubule pausing followed by growth, thus effectively preventing microtubule depolymerization. This effect of CSPP1 on microtubule behavior strikingly resembles that of microtubule-stabilizing compounds, taxanes and epothilones, which also preferentially accumulate at growing microtubule ends in pre-catastrophe state, causing their stabilization and pausing followed by polymerization (Rai et al., 2020). Since taxanes bind to microtubules from the luminal side, we hypothesized that the same would be true for CSPP1. We investigated CSPP1 localization using cryo-electron tomography (cryo-ET) and MINFLUX microscopy and found that CSPP1 is a microtubule inner protein (MIP). In line with this finding, we observed that in addition to localizing at growing microtubule ends, CSPP1 also efficiently binds to sites where microtubule lattices are damaged. Furthermore, deletion mapping showed that CSPP1 contains separate domains responsible for microtubule rescue and stabilization and for growth inhibition. Altogether, our findings reveal how microtubule dynamics can be controlled from the luminal side. These data have important implications for understanding how highly stable microtubule populations, such as those in ciliary axonemes, are generated and maintained.

## Results

### CSPP1 suppresses catastrophes by binding to polymerizing ends where it induces pausing

CSPP1 contains several predicted helical domains interspersed with regions of unknown structure and is represented by two isoforms, the long isoform CSPP-L and a shorter isoform (termed here CSPP-S), which lacks 294 amino acids at N-terminus and contains an internal deletion of 52 amino acids (Fig. 1A, (Frikstad et al., 2019; Patzke et al., 2006)). In our initial analysis, we focused on the CSPP-L isoform. To get insight into the autonomous effects of CSPP-L on microtubule dynamics, we have purified it from HEK293 cells (Fig. S1A). Mass spectrometry-based analysis demonstrated that CSPP-L preparations contained no other known regulators of microtubule dynamics (Fig. S1B). We used purified GFP-CSPP-L to perform in vitro assays where microtubules grown from GMPCPP-stabilized seeds were observed by Total Internal Reflection Fluorescence microscopy (Fig. S1C)(Aher et al., 2018; Bieling et al., 2007). In the presence of tubulin alone, microtubules regularly switched from growth to shortening that proceeded all the way back to the seed. However, the addition of 10 nM CSPP-L suppressed shrinkage and led to frequent pausing of microtubule plus ends, while their growth rate was slightly reduced (Fig. 1B, C, E, F; Fig. S1D, E; Video S1). A similar effect was observed when we included in the assay mCherry-EB3, a marker of growing microtubule ends, which by itself increases microtubule growth rate and promotes catastrophes (Fig. 1D-F; Fig. S1E)(Komarova et al., 2009). In our in vitro assays, CSPP-L also bound to growing microtubule minus ends and strongly accumulated along the lattice formed by minus-end polymerization (Fig. 1B-D; Fig. S1E). However, in cells, this protein normally acts at the distal tip of the cilium which contains microtubule plus ends, and therefore we have not investigated the effects of CSPP-L on microtubule minus-end dynamics. CSPP-L binding was always initiated close to the growing microtubule end, and after binding, CSPP-L showed little lateral diffusion along microtubules, so that CSPP-L binding zones remained well-confined (Fig. 1C, D). The low lateral mobility of CSPP1 was confirmed by spiking experiments where 0.5 nM GFP-CSPP-L was combined with 9.5 nM mCherry- CSPP-L (Fig. S1F). When CSPP-L concentration was increased, the zones of CSPP-L accumulation coincided with longer and more frequent microtubule pausing events (Fig. 1D-F). CSPP-L-induced pausing was almost always (in ∼95% of the cases) followed by growth and not by shrinkage, and at CSPP-L concentrations exceeding 5 nM, very little microtubule depolymerization was observed (Fig. 1B-F; Fig. S1E). At low, 0.5 nM concentration of CSPP-L, long microtubule depolymerization episodes were still present, but zones of CSPP-L accumulation triggered microtubule rescues (Fig. 1D, F).

**Figure 1:**
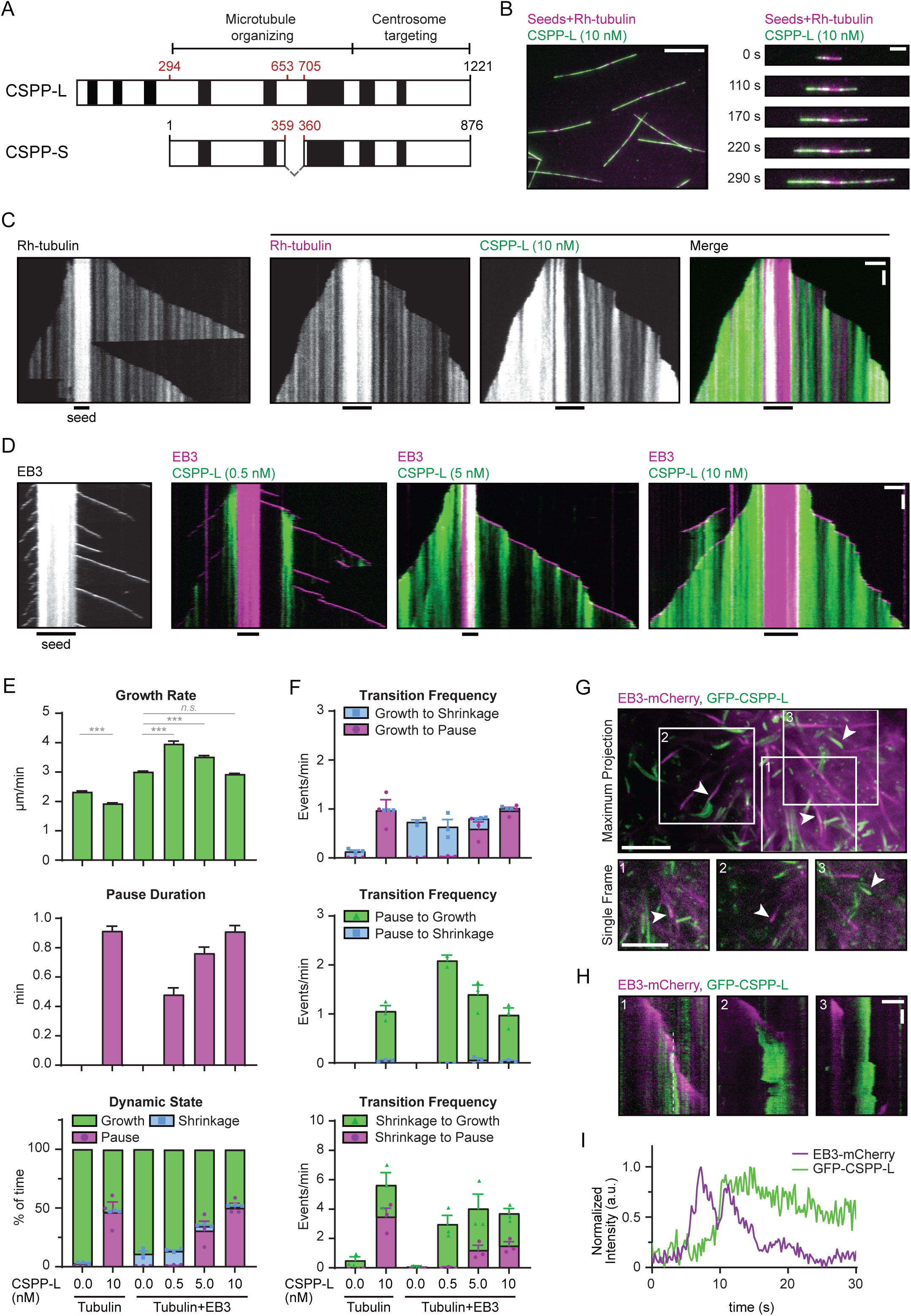
**CSPP1 suppresses catastrophes by binding to polymerizing ends where it induces pausing** A. Schematic representation of the two isoforms expressed by the *CSPP1* gene in mammals. Black boxes represent α-helical domains larger than 20 amino acids predicted by AlphaFold. B. Field of view (left, scale bar 10 µm) and time-lapse images (right, scale bar 3 µm) illustrating microtubule growth from GMPCPP stabilized microtubule seeds in the presence of 15 µM tubulin supplemented with 3% rhodamine-labelled tubulin and 10 nM GFP-CSPP-L. C,D. Kymographs illustrating microtubule growth either with rhodamine-tubulin (C) or mCherry- EB3 (D), supplemented, where indicated, with the indicated concentrations of GFP-CSPP-L. Scale bars, 2 μm (horizontal) and 60 s (vertical). E, F. Parameters of microtubule plus end dynamics in the presence of rhodamine-tubulin alone or together with 20 nM mCherry-EB3 in combination with the indicated GFP-CSPP-L concentrations (from kymographs as shown in (C, D). Events were classified as pauses when the pause duration was longer than 20 s. Number of growth events, pauses and microtubules analyzed (E); tubulin alone, n= 394, 0, 110; tubulin with 10 nM CSPP-L, n= 596, 481, 78; EB3 alone, n= 514, 0, 53; EB3 with 0.5 nM CSPP-L, n=731, 10, 44; EB3 with 5 nM CSPP-L, n=564, 241, 47; EB3 with 10 nM CSPP-L, n=476, 518, 89. Number of transition events analyzed (F): tubulin alone, n= 194, 0, 0, 0, 15, 0; tubulin with 10 nM CSPP-L, n= 0, 443, 410, 25, 7, 17; EB3 alone, n= 461, 0, 0, 0, 4, 0; EB3 with 0.5 nM CSPP-L, n=309, 8, 10, 0, 216, 2; EB3 with 5 nM CSPP-L, n=75, 209, 224, 9, 57, 27; EB3 with 10 nM CSPP-L, n=24, 465, 455, 22, 25, 21. Single data points represent averages of three independent experiments. Error bars represent SEM. ***, p< 0.001; n.s., not significant; Kruskal-Wallis test followed by Dunńs post-test. G. Maximum projection (top) and enlarged single frame images (bottom) of a COS-7 cell overexpressing GFP-CSPP-L and EB3-mCherry, imaged by TIRF microscopy. Scale bars, 5 µm. H. Kymographs of the events indicated with arrowheads in (G). Scale bars, 2 μm (horizontal) and 4 s (vertical). I. Normalized intensity graphs of EB3-mCherry and GFP-CSPP-L along the white dashed line in (H). See also Figure S1 and Videos S1 and S2.

Next, we investigated the behavior of GFP-CSPP-L in COS-7 cells. Endogenous CSPP1 in these cells is only localized to centrioles and centriolar satellites but not to cytoplasmic microtubules (Fig. S1G). Similar to what we observed in vitro, overexpressed GFP-CSPP-L formed accumulations along microtubules (Fig. 1G; Fig. S1H; Video S2). Elevated levels of CSPP-L led to an increase in microtubule acetylation (Fig. S1H, I), a hallmark of microtubule stabilization (Magiera et al., 2018). Moreover, the number of microtubule plus ends labeled with EB1, a marker of growing microtubule ends (Mimori-Kiyosue et al., 2000), was strongly reduced (Fig. S1H, J), indicating that microtubule dynamics was suppressed. Live cell imaging in COS-7 cells co-expressing GFP-CSPP-L and EB3- mCherry showed that CSPP-L bound to growing, EB3-positive microtubule ends upon EB3 signal reduction, and CSPP-L accumulation led to microtubule pausing (Fig. 1G-I). Microtubules could regrow from CSPP-L accumulations (Fig. 1G, H, box 1) or stay paused for longer periods of time (Fig. 1G, H, box 2), and many pausing microtubule ends strongly labeled with CSPP-L were observed throughout the cell (Fig. 1G, H, box 3, Video S2). We conclude that CSPP-L binds to growing microtubule ends, prevents their shrinkage and induces pausing both in vitro and in cells.

### CSPP1 binds to pre-catastrophe microtubule ends, resembling taxane behavior

Formation of confined accumulation zones that initiate at growing microtubule ends and prevent microtubule shrinkage makes the dynamic behavior of CSPP-L strikingly similar to that we have recently described for taxanes (Rai et al., 2020). To determine if CSPP-L and taxanes recognize the same features of microtubules, we have tested whether fluorescently labelled taxane Fchitax-3 colocalized with CSPP-L and found that this was indeed the case (Fig. 2A, B). Over time, the intensity of both Fchitax-3 and CSPP-L first increased and then decreased in a similar way (Fig. 2A, C). Measurements of fluorescence intensity of 10 nM CSPP-L within accumulation zones, performed as described previously (Rai et al., 2020), indicated that on average, one CSPP-L molecule was bound per 8 nm of microtubule length (corresponding to the length of one layer of α/β-tubulin dimers) (Fig. 2D), indicating that the binding sites are likely not saturated in these conditions.

**Figure 2:**
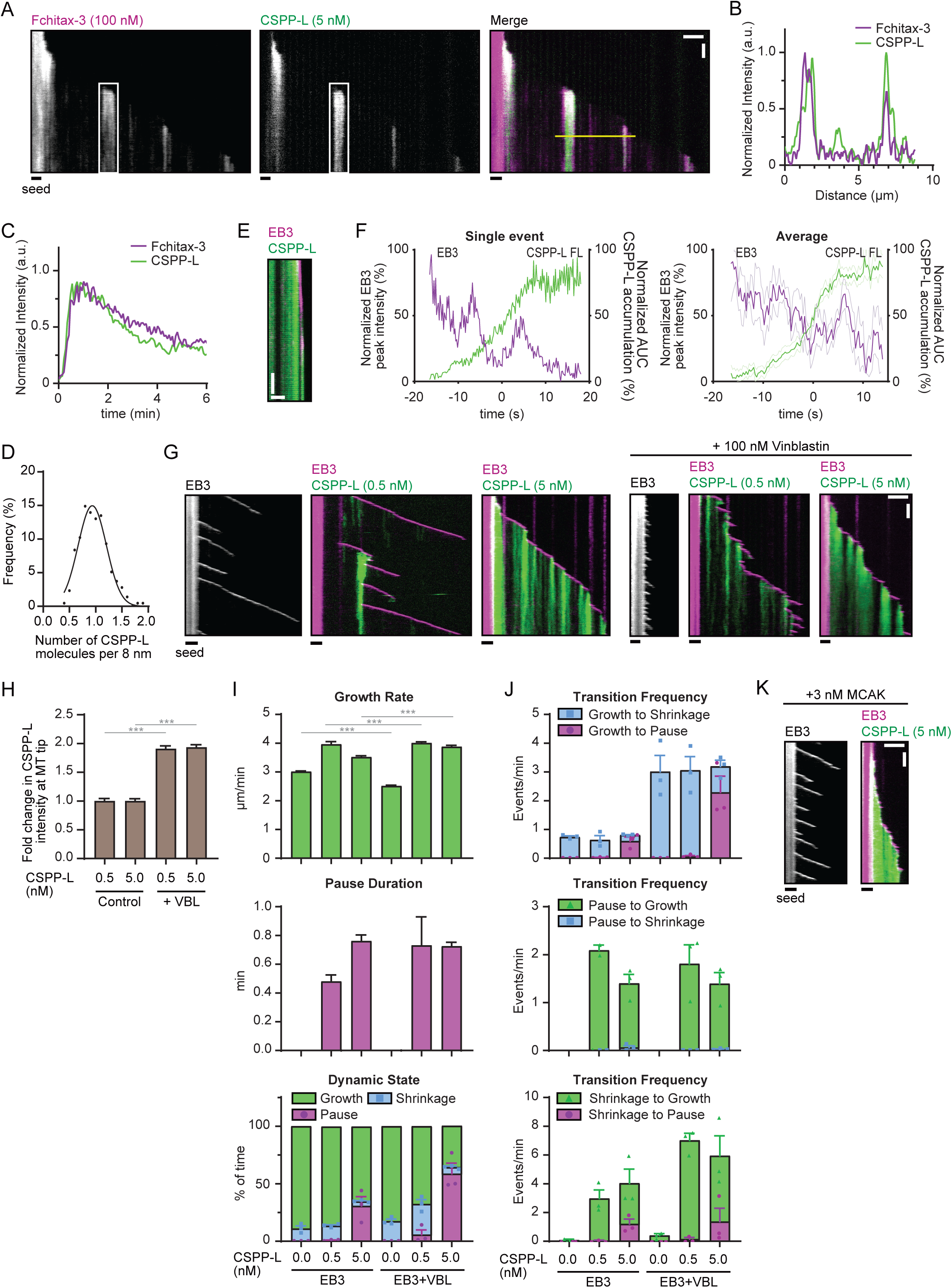
**CSPP1 binds to pre-catastrophe microtubule ends, resembling taxane behaviour** A. Kymographs of microtubule growth with 100 nM Fchitax-3 together with 5 nM mCherry-CSPP- L in presence of 20 nM dark EB3. Scale bars, 2 μm (horizontal) and 60 s (vertical). B. Normalized intensity graph of Fchitax-3 and GFP-CSPP-L along the yellow line in (A). C. Normalized intensity graph of Fchitax-3 and GFP-CSPP-L within the white box in (A). D. Quantification of the number of GFP-CSPP-L molecules per 8 nm microtubules. The integrated intensity of one GFP-CSPP-L accumulation in an in vitro assay was divided by the average intensity of single GFP monomers in a separate chamber on the same coverslip and subsequently normalized to 8 nm accumulation length. Number of GFP-CSPP-L accumulations n=215. E. Kymograph illustrating microtubule growth in the presence of 20 nM mCherry-EB3 together with 10 nM GFP-CSPP-L. Scale bars, 2 μm (horizontal) and 5 s (vertical). F. Time plot of the normalized maximum intensity profile of a single mCherry-EB3 comet and the normalized area under the curve (AUC) of a single GFP-CSPP-L accumulation (left) and averaged EB3 and GFP-CSPP-L profiles, normalized and aligned using half-maximum effective intensity values from Hill equation fits as reference points (right) (from kymographs as shown in E). Dashed lines represent SEM. Number of events analyzed, n= 12 from two independent experiments. G. Kymographs of microtubule growth with mCherry-EB3 alone or together with indicated concentrations of GFP-CSPP-L in presence or absence of 100 nM Vinblastin (VBL). Scale bars, 2 μm (horizontal) and 60 s (vertical). H. Quantification of the mean GFP-CSPP-L intensity at the microtubule tip per growth event. The average mean intensity of GFP-CSPP-L in presence of 100 nM Vinblastin was normalized to the average mean intensity in absence of Vinblastin. Number of growth events analyzed; 0.5 nM GFP- CSPP-L control, n=474; 5 nM GFP-CSPP-L control, n=598; 0.5 nM GFP-CSPP-L with Vinblastin, n=1363; 5 nM GFP-CSPP-L with Vinblastin, n=897. I,J. Parameters of microtubule plus end dynamics in the presence of 20 nM mCherry-EB3 together with the indicated GFP-CSPP-L concentrations (from kymographs as shown in G). Events were classified as pauses when the pause duration was longer than 20 s. Number of growth events, pauses and microtubules analyzed (I); EB3 alone, n= 514, 0, 53; EB3 with 0.5 nM CSPP-L, n=731, 10, 44; EB3 with 5 nM CSPP-L, n=564, 241, 47; EB3 with Vinblastin, n=915, 0, 54; EB3 with 0.5 nM CSPP- L and Vinblastin, n=1204, 33, 40; EB3 with 5 nM CSPP-L and Vinblastin, n=632, 408, 47. Number of transition events analyzed (J): EB3 alone, n= 461, 0, 0, 0, 4, 0; EB3 with 0.5 nM CSPP-L, n=309, 8, 10, 0, 216, 2; EB3 with 5 nM CSPP-L, n=75, 209, 224, 9, 57, 27; EB3 with Vinblastin, n=162, 0, 0, 0, 33, 0; EB3 with 0.5 nM CSPP-L and Vinblastin, n=1079, 19, 31, 0, 1002, 14; EB3 with 5 nM CSPP-L and Vinblastin, n=147, 372, 386, 7, 127, 27. Single data points represent averages of three independent experiments. Data for conditions without Vinblastin is the same as in Fig. 1E,F. K. Kymographs illustrating microtubule growth with mCherry-EB3 alone or together with 5 nM GFP-CSPP-L in presence of 3 nM MCAK. Scale bars, 2 μm (horizontal) and 60 s (vertical). For all plots. Error bars represent SEM. ***, p< 0.001; n.s., Kruskal-Wallis test followed by Dunńs post-test.

Since our previous work has demonstrated that binding of Fchitax-3 is triggered by perturbed microtubule growth and occurs when microtubules enter a pre-catastrophe state manifested by the loss of GTP cap and reduced EB3 binding (Rai et al., 2020), we tested whether the same is true for CSPP-L. Indeed, periods of strong CSPP-L accumulation always initiated a few seconds after EB3 signal started to diminish (Fig. 2E, F), and a similar CSPP-L accumulation pattern was observed in cells (Fig. 1G-I). To support our interpretation that perturbed microtubule growth triggers CSPP-L binding, we supplemented the assay with 100 nM vinblastine, which promotes frequent catastrophes at low concentrations in the presence of EB3 (Mohan et al., 2013). Catastrophes indeed became much more frequent, and this resulted in the increased number of CSPP-L accumulation zones, leading to a higher overall binding of the protein along microtubules (Fig. 2G, H, J). Similar to the conditions without vinblastine, 0.5 nM CSPP-L did not block depolymerization completely but induced formation of rescue sites, whereas 5 nM CSPP-L induced more frequent pausing episodes followed by re-growth (Fig. 2G, I, J). In the presence of the kinesin-13 MCAK, which also triggers frequent catastrophes in the presence of EB3 (Montenegro Gouveia et al., 2010), enhancement of CSPP-L accumulation along microtubules was observed as well (Fig. 2K). We conclude that similar to taxanes, CSPP-L strongly accumulates at microtubule ends that undergo a growth perturbation, inhibits both their growth and shortening and gradually dissociates when microtubule growth resumes.

### Separate CSPP1 domains control the balance between microtubule polymerization and depolymerization

Next, we examined which CSPP1 domains are responsible for its effects on microtubule dynamics. As a starting point for deletion mapping of CSPP1, we used structure predictions made by a recently developed neural network AlphaFold (Jumper et al., 2021; Varadi et al., 2022). As mentioned previously, CSPP-L contains several putative α-helical domains (H1-8) interspersed with regions of unknown structure (L1-L7) (Fig. 3A). Compared to the previously performed analyses of CSPP1 sequence, this prediction suggested presence of two additional α-helical regions, H4 and H8, in the middle and C-terminal part of the protein. Based on the predicted domains, we generated various GFP-tagged fragments of CSPP1 (Fig. 3A) and tested them in the in vitro assays. The short isoform of CSPP1, CSPP-S, behaved similarly to CSPP-L, though at 10 nM it was less efficient at preventing microtubule depolymerization and could also occasionally block microtubule outgrowth from the seed, whereas we have never observed this effect with CSPP-L (Fig. 3A; S2A). Next, we focused on the middle part of CSPP1, because previous work has identified it as the microtubule-organizing region (Frikstad et al., 2019; Patzke et al., 2006) (MTOR, Fig. 3A; S2B, C). The MTOR region derived from the CSPP-L isoform displayed local accumulations along microtubules and prevented catastrophes at 10 nM, but did not cause long pauses, even at 40 nM concentration (Fig. S2B). The MTOR version with the internal deletion present in CSPP-S showed little microtubule binding at 10 nM, but the binding became visible at 40 nM and was accompanied by frequent pauses, followed by either growth or shrinkage (Fig. 3A; S2C). Further deletion mapping at the C-terminus of the MTOR domain (the construct H4+L4+H5) showed that the helical domain H6 with the preceding linker L5 was not essential for microtubule binding or rescue activity but was needed to trigger pausing (Fig. 3A; S2D). An even shorter truncation mutant, which also lacked helical domain H5 (H4+L4) displayed only a very weak binding to microtubules (Fig. 3A; S2E). However, the affinity of this fragment for microtubules was increased by linking it to the leucine zipper dimerization domain of GCN4 (H4+L4+LZ) (Fig. 3A, S2F). Therefore, from this point onwards we termed helical domain H5 the dimerization domain (DD). Linking the newly identified short α-helical domain H4 directly to the leucine zipper through a short flexible linker yielded again a construct that weakly bound to microtubules but did not induce rescues, even at concentrations up to 300 nM (H4+LZ; Fig. 3A-D; Fig. S2G). Extension of H4 with a part of linker L4 (amino acids 375-453; a protein fragment we termed MTB, for “microtubule-binding”), fused to the leucine zipper, resulted in a construct which was sufficient for microtubule binding and rescue induction (MTB+LZ; Fig. 3A-D; Fig. S2H). Microtubule binding of CSPP1 thus depends on a short region, which is predicted to be α-helical, and is augmented by dimerization and additional regions distributed throughout the CSPP1 molecule, including the region missing in the CSPP-S isoform.

**Figure 3:**
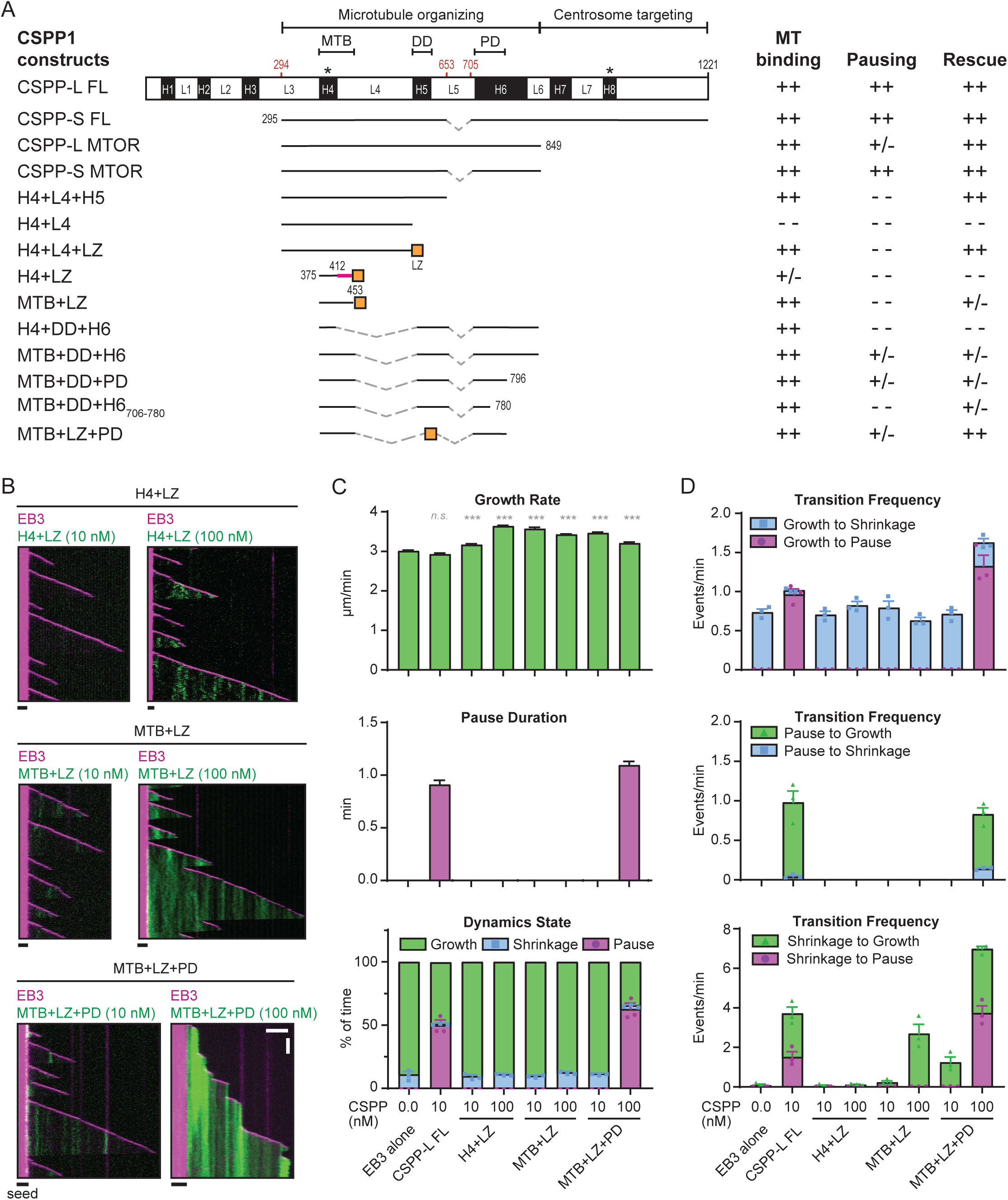
**Separate CSPP1 domains control the balance between microtubule polymerization and depolymerization** A. Schematic representation of the different CSPP1 constructs used and a summary of their binding to microtubules and their effects on microtubule dynamics. Black boxes represent α-helical domains larger than 20 amino acids predicted by AlphaFold; asterisks indicate previously unidentified helices. ++: frequently observed at protein below 40 nM; +/-: occasionally observed at protein concentrations below 40 nM and/or frequently observed at protein concentrations up to 100 nM; - -: observed infrequently or not observed at all even at protein concentrations higher than 100 nM. B. Kymographs illustrating microtubule growth with 20 nM mCherry-EB3 alone or together with 10 or 100 nM of the indicated GFP-CSPP1 constructs. Scale bars, 2 μm (horizontal) and 60 s (vertical). C,D. Parameters of microtubule plus end dynamics in the presence of 20 nM mCherry-EB3 together with 10 or 100 nM of the indicated GFP-CSPP1 constructs (from kymographs as shown in B). Events were classified as pauses when the pause duration was longer than 20 s. Number of growth events, pauses and microtubules analyzed (C); EB3 alone, n= 514, 0, 53; EB3 with 10 nM CSPP-L, n=24, 465, 455, 22, 25, 21; EB3 with 10 nM H4+LZ, n=855, 0, 87; EB3 with 100 nM H4+LZ, n=987, 0, 103; EB3 with 10 nM MTB+LZ, n=1006, 0, 109; EB3 with 100 nM MTB+LZ, n=1206, 0, 139; EB3 with 10 nM MTB+LZ+PD, n=934, 0, 104; EB3 with 100 nM MTB+LZ+PD, n=776, 707, 123. Number of transition events analyzed (D): EB3 alone, n= 461, 0, 0, 0, 4, 0; EB3 ith 10 nM CSPP-L, n=24, 465, 455, 22, 25, 21; EB3 with 10 nM H4+LZ, n=751, 0, 0, 0, 3, 0; EB3 with 100 nM H4+LZ, n=889, 0, 0, 0, 15, 0; EB3 with 10 nM MTB+LZ, n= 902, 0, 0, 0, 26, 0; EB3 with 100 nM MTB+LZ, n= 1035, 0, 0, 0, 582; EB3 with 10 nM MTB+LZ+PD, n=797, 0, 0, 0, 191, 0; EB3 with 100 nM MTB+LZ+PD, n=126, 545, 520, 105, 107, 121. Single data points represent averages of three independent experiments. Error bars represent SEM. ***, p< 0.001; n.s., not significant; Kruskal-Wallis test followed by Dunńs post-test. Data for EB3 alone and EB3 with 10 nM CSPP-L is the same as in Fig. 1E,F. See also Figure S2.

Importantly, all CSPP1 fragments lacking the domain H6 did not cause microtubule pausing, suggesting that H6 could be responsible for pause induction. To test this idea, we first directly fused the DD and H6 domains to H4 (H4+DD+H6). Already at 40 nM concentration, this construct strongly inhibited microtubule outgrowth from the seed and induced catastrophes (Fig. 3A; S2I). Attaching the DD and H6 domains to MTB (MTB+DD+H6) resulted in a construct that could induce microtubule pausing and inhibit depolymerization at 100 nM, whereas at lower concentrations (40 nM), it showed occasional rescues but no pauses (Fig. 3A; S2J). To determine which part of H6 is responsible for inhibiting microtubule growth, we truncated it at the C-terminus and found that MTB+DD+H6_706-796_ but not a shorter version, MTB+DD+H6_706-780_, still triggered pausing and inhibited microtubule shrinkage when fused to MTB and H5 (Fig. 3A, Fig. S2K, L). We therefore termed H6_706-796_ the pausing domain (PD). Swapping DD within this construct for the leucine zipper (MTB+LZ+PD) yielded a construct with similar properties (Fig. 3A-D; Fig. S2M), confirming that DD primarily acts as a dimerization domain.

Next, we compared the impact of truncated CSPP1 constructs with that of GFP-CSPP-L on microtubules in COS-7 cells. GFP-MTB+LZ+PD, GFP-MTB+LZ and GFP-H4+LZ localized to microtubules in interphase cells. However, compared to CSPP-L, the shorter constructs were less potent in inducing microtubule acetylation and reducing the number of EB1 comets, indicating that they are less efficient in stabilizing microtubules (Fig. S2N-Q; control in Fig. S1H-J). Altogether, we conclude that CSPP1 has multiple regions contributing to microtubule binding, but the minimal construct that reproduces the major effects of CSPP-L on microtubule dynamics is MTB+LZ+PD. These effects appear to depend on the interplay between two separate activities, residing in two predicted helical regions: microtubule binding and stabilization by the MTB and the growth-inhibiting activity of the truncated α-helical domain H6, the PD.

### CSPP1 binds to microtubule lumen

As described above, the behavior and effect of CSPP1 on dynamic microtubules resembles that of taxanes. Taxanes are known to bind to the microtubule lumen (reviewed in (Steinmetz and Prota, 2018)), and therefore, we set up cryo-ET experiments to investigate whether CSPP1 is a MIP. Using a previously established experimental design (Ogunmolu et al., 2021), we polymerized dynamic microtubules from GMPCPP-stabilized seeds in presence or absence of 10 nM CSPP-L, with or without 250 nM vinblastine and vitrified them on EM grids. To increase the signal-to-noise ratio in the reconstructed tomograms, we used the cryoCARE denoising method (Buchholz et al., 2019). Microtubules polymerized in presence of CSPP-L frequently contained luminal densities, which were absent in CSPP-L-free samples (Fig. 4A, B; Fig. S3A). Presence of vinblastine resulted in higher percentage of microtubules containing MIP particles: 68% ± 22% (mean ± SD), compared to 35% ± 11% in absence of vinblastine (*p* < 10^-4^, Fig. 4A, B). We did not observe CSPP-L densities outside of microtubule lumen.

**Figure 4:**
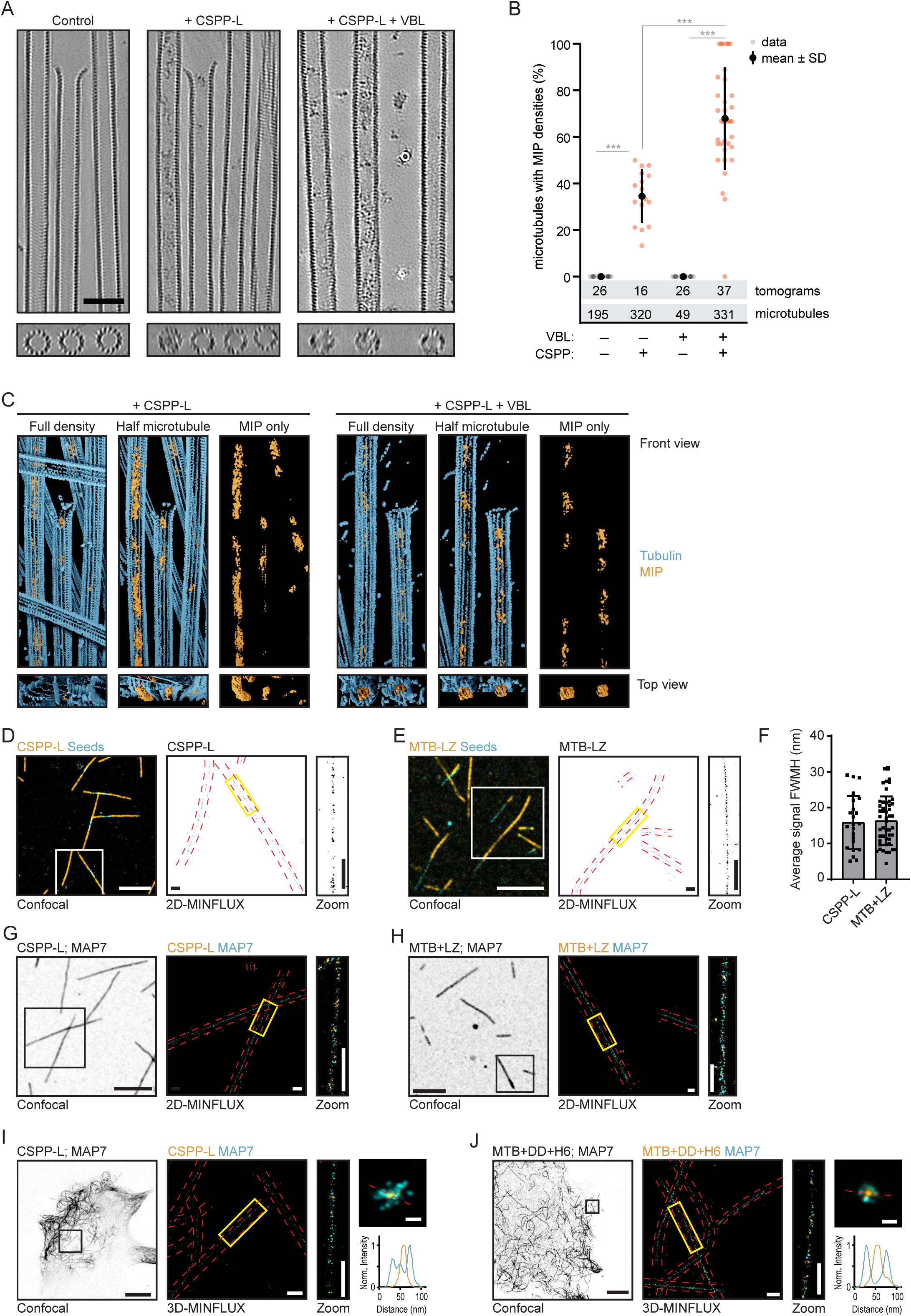
**CSPP1 binds to microtubule lumen.** A. Denoised tomograms of dynamic microtubules polymerized from GMPCPP-stabilized seeds in the presence or absence of 10 nM GFP-CSPP-L, with or without 250 nM Vinblastine vitrified on EM grids. Scale bar, 50 nm. B. Quantification of the percentage of microtubules containing luminal densities from total microtubules (from tomograms as shown in A). Orange and grey dots (single data points, tomograms), black circle (mean), SD (error bars). Number of microtubules and tomograms are displayed in the graph. ***, p< 0.001, Mann-Whitney test. Analysis from two experiments. C. Reconstituted images from automated segmentation of denoised tomograms as in (A). D-E. Single color 2D-MINFLUX measurements of in vitro reconstituted microtubules polymerized in the presence of SNAP-CSPP-L (D) or SNAP-MTB-LZ (E). Images were rendered with 1 nm voxel size for visualization. White boxes in confocal image indicate the region shown in the rendered 2D- MINFLUX image, yellow boxes in the 2D-MINFLUX image indicate the region of the zoom, red dashed lines represent the microtubule outline from the confocal image. Scale bars; 5 μm (Confocal image), 500 nm (2D-MINFLUX image and Zoom). F. Quantification of the fitted, Full-Width-Half-Maximum values per microtubule (from 2D- MINFLUX images as shown in D and E). Single data points are shown. SD (error bars). Number of measured microtubules; CSPP-L, n = 23; MTB+LZ, n = 49. Analysis from three experiments. G-H. Dual color 2D MINFLUX measurements of in vitro reconstituted microtubules polymerized in the presence of GFP-MAP7 N-terminus together with SNAP-CSPP-L (G) or SNAP-MTB-LZ (H). Images were rendered with 4 nm voxel size for visualization. Panel representation as in (D-E). I-J. Dual color 3D-MINFLUX measurements of COS-7 cells overexpressing GFP-MAP7 plus SNAP- CSPP-L (I) or SNAP-MTB-DD-H6 (J). Images were rendered with 4 nm voxel size for visualization. Black boxes in confocal image indicate the region shown in the rendered 3D-MINFLUX image, yellow boxes in the 3D-MINFLUX image indicate the region of the zoom, red dashed lines represent the microtubule outline from the confocal image. Top right image shows a maximum intensity projection of the cross section of the microtubule over 400 nm (CSPP-L) or 800 nm (MTB+DD+H6). The red dashed line there indicates the line scan related to the bottom right graph. Scale bars; 10 μm (Confocal image), 500 nm (2D-MINFLUX image and Zoom), 50 nm (maximum intensity projection image). See also Figure S3 and Videos S3 and S4.

We further used automated segmentation of denoised tomograms (Chen et al., 2017) to get a better understanding of the intraluminal particles. CSPP-L particles appeared quite disordered, and could either block the microtubule lumen completely, or only partially (Fig. 4C, Video S3). They were occupying variable length of the microtubule lumen, preventing further analysis of their structure. Some CSPP-L particles were bound close to the terminal flare of tubulin protofilaments, but we never observed them binding to tapered microtubule ends or other incomplete microtubule lattices.

Next, we aimed to confirm that the densities inside microtubules we observed with Cryo-ET indeed represent CSPP-L and determine the localization of shorter CSPP1 fragments. Since the latter would be difficult to achieve by cryo-ET due to the small protein size, we turned to MINFLUX microscopy, which allows localization of individual fluorophores with very high spatial resolution, as was demonstrated by the separation of e.g. two fluorophores as close as 6 nm from each other (Balzarotti et al., 2017; Gwosch et al., 2020). The localization resolution of MINFLUX would allow us to determine whether the CSPP1 fragments localize inside or outside 25-nm-wide microtubule filaments. For 2D MINFLUX measurements, we used fixed microtubules that were grown in vitro in the presence of SNAP-tagged CSPP1 or its fragments. We first performed measurements for CSPP-L, the same protein we used for cryo-ET. We determined the Full-Width-Half-Maximum (FWHM) values of the measured localizations and found the diameter CSPP-L signal was 15.87 ± 7.47 nm (mean ± SD) (Fig. 4D, F; Fig. S3B), clearly indicating it is inside microtubule lumen. This is supported by the determined localization precision in x and y of the MINFLUX measurements, as these values are 3.7 and 3.2 nm, respectively (Fig. S3C). The smallest CSPP1 fragment binding to microtubules, H4+LZ, gave too much background signal to allow meaningful measurements. However, the smallest CSPP1 construct affecting microtubule dynamics in vitro, MTB+LZ (Fig. 3A- C) gave a signal diameter of 16.35 ± 6.80 nm (mean ± SD), also indicating that it is located inside the microtubule lumen (Fig. 4E, F; Fig. S3B). To validate that the signal width is due to the protein being inside the microtubule lumen, we performed the same assays with GFP-labeled N-terminal, microtubule-binding part of MAP7, a protein known to bind to microtubule exterior (Ferro et al., 2022), which was also added during microtubule polymerization. Dual color 2D MINFLUX measurements showed that both CSPP-L and MTB+LZ were surrounded by the MAP7 signal (Fig. 4G, H; Fig. S3B). The distinct localization of CSPP1 and MAP7, a label for the outer microtubule surface, together with the FWHM analysis indicate that CSPP-L and its short microtubule-binding fragment indeed localize to the lumen of in vitro reconstituted microtubules.

Next, we tested whether CSPP1 fragments localizes inside microtubules in mammalian cells. We overexpressed SNAP-CSPP-L and GFP-MAP7 in COS-7 cells, fixed them and stained them for SNAP and GFP. Even though the labeling density of CSPP-L was very sparse, the maximum intensity projection over the cross section of the microtubule showed a ring of MAP7 signal surrounding CSPP-L (Fig. 4I; Fig. S3D). This was even more striking when we overexpressed the smaller SNAP- MTB+DD+H6 fragment together with GFP-MAP7 (Fig. 4J, Fig. S3D, E; Video S4). Acquisition of MINFLUX images for even shorter CSPP1 fragments were impeded by the high cytosolic background due to the presence of a significant pool of microtubule-unbound proteins. Taken together, the data obtained in vitro and in cells support the intraluminal localization of CSPP1 and indicate that the short MTB domain is sufficient for this localization.

### CSPP1 efficiently binds to sites where microtubule lattices are damaged

Since CSPP1 is a MIP, we next examined whether it can bind to sites of lattice damage, which would provide access to microtubule lumen. First, we compared the binding of CSPP1 to Taxol-stabilized microtubules, which are known to acquire extensive lattice defects when incubated without soluble tubulin (Aher et al., 2020; Arnal and Wade, 1995), to more stable GMPCPP-bound microtubules. In the absence of soluble tubulin, CSPP-L gradually accumulated at discrete sites on both types of microtubules, but the binding to Taxol-stabilized microtubules was stronger and faster (Fig. 5A, B). Next, we induced local damage of GMPCPP-stabilized microtubules using illumination with a pulsed 532 nm laser, as described previously (Aher et al., 2020). We chose microtubule regions where no prior CSPP-L signal was present and selected for analysis only the microtubules that were not fully severed during laser illumination. We observed strong accumulation of CSPP-L at the illuminated sites, whereas the CSPP-L signal was relatively stable within the same time period at the sites that were not damaged with the laser (Fig. 5C-E).

**Figure 5:**
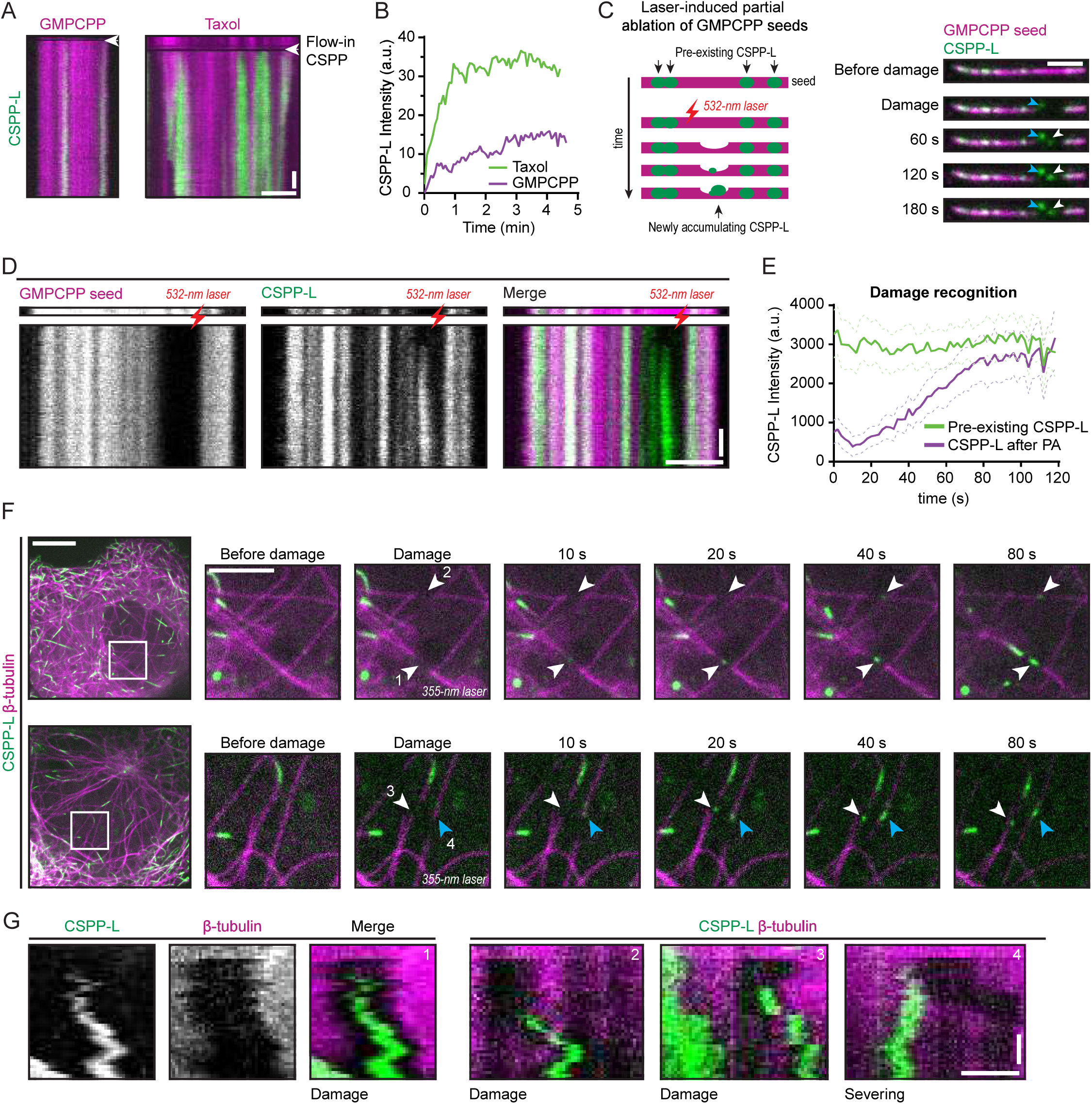
**CSPP1 binds to sites where microtubule lattices are damaged** A. Kymographs of GMPCPP- (left) and Taxol-stabilized (right) microtubule seeds. 5 nM GFP-CSPP- L was flushed in during acquisition, in absence of free taxol or tubulin. Scale bars, 2 μm (horizontal) and 30 s (vertical). B. GFP-CSPP-L intensity profile of developing accumulation after flow-in of experiments done in (A). C. Schematic representation (left) and time lapse images (right) of laser damage of a GMPCPP- stabilized microtubule seed at regions with no prior GFP-CSPP-L accumulation. The microtubule region illuminated with the 532 nm pulsed laser is highlighted by a white arrowhead. The blue arrowhead indicates the damage inflicted on the coverslip. Scale bar 2 µm. D. Kymograph corresponding to time lapse images shown in (C). The laser illuminated microtubule region is highlighted by a red lightning bolt. Scale bars, 2 μm (horizontal) and 30 s (vertical). E. Averaged GFP-CSPP-L intensity profiles of after photodamage (from kymographs as shown in D). Plots were aligned using half-maximum effective intensity values from nonlinear regression fits as reference points. Dashed lines represent SEM. Number of events analysed, n= 15 from three independent experiments. F. Time lapse images of photodamage experiments in COS-7 cells overexpressing GFP-CSPP-L and β-tubulin-mCherry. Arrowheads indicate the events where microtubules were damaged (white) or severed (blue). Imaging was performed using spinning disk microscopy and photodamage was induced with a 355 nm laser. Scale bars 10 µm (left) and 4 µm (zoom). G. Kymographs of the events shown in (F). Scale bars, 1 μm (horizontal) and 20 s (vertical). See also Video S5.

Finally, we examined whether CSPP-L can recognize sites of microtubule damage in cells by performing laser microsurgery in COS-7 cells co-expressing GFP-CSPP-L and β-tubulin-mCherry. We damaged single microtubules in the z-plane just below the nucleus by local illumination with a 355 nm laser and observed CSPP-L accumulations forming at the illuminated positions (Fig. 5F, G; Video S5). It was more difficult to introduce local microtubule damage by laser microsurgery in cells than in vitro, because the intensity of the laser beam varied with microtubule positions in the z-plane, so the degree of the photodamage was difficult to predict. For the analysis, we only considered events where new CSPP-L signal appeared at the position where the microtubule intensity was reduced after laser illumination. To distinguish partial damage from complete severing, we focused on the events where the illuminated microtubule was visible on both sides of the newly formed CSPP-L accumulation and where both microtubule parts moved synchronously with the photobleached region (Fig. 5F, G). The average time between laser illumination and the appearance of CSPP-L signal was 21 ± 13 s and the size of the CSPP-L accumulation 564 ± 157 nm (mean ± SD, n = 83). Thus, CSPP1 can bind to damaged microtubule lattices in vitro and in cells.

### CSPP1 stabilizes damaged microtubules and promotes lattice integrity

To determine whether CSPP1 can stabilize damaged microtubules, we again used Taxol-stabilized microtubules. Binding of CSPP-L to Taxol-stabilized microtubules was suppressed by the presence of free Taxol, which can prevent microtubule disassembly and erosion (Fig. 6A, B; Fig. S4A). In the absence of Taxol in solution, Taxol-stabilized microtubules gradually depolymerized (Fig. 6A). To quantify the effects of Taxol, CSPP-L and free tubulin on microtubule stability, we determined the percentage of Taxol-stabilized seeds surviving after 5 min (Figure 6C). CSPP-L could slow down though not block microtubule depolymerization in a concentration-dependent manner (Fig. 6A, C). The addition of free Taxol to these assays stabilized the microtubules completely, but when CSPP-L was also present, stabilization was slightly reduced, suggesting a potential competition between Taxol and CSPP1 for microtubule binding. The addition of low concentrations of free tubulin (2-5 µM) in the absence of free Taxol had a very mild stabilizing effect in these assays, but in the presence of CSPP-L, complete microtubule stabilization was observed already at 2 µM tubulin (Fig. 6A, C). At 5 µM tubulin, CSPP-L even facilitated new microtubule lattice outgrowth (Fig. 6A), indicating that it might lower the tubulin concentration threshold for templated microtubule polymerization, as previously observed with some other microtubule regulators (Aher et al., 2018; Wieczorek et al., 2015). To confirm this conclusion, we repeated the assays with GMPCPP-stabilized microtubule seeds and found that CSPP-L strongly increased the frequency of microtubule outgrowth from seeds at 5 µM tubulin (Fig. 6D; Fig. S4B). Interestingly, CSPP-L intensity along the newly formed microtubule lattice was much higher when microtubules were grown in 5 µM tubulin compared to 15 µM tubulin (Fig. 6E, F). This suggests that CSPP-L binds to microtubules more efficiently when they grow slowly, either due to a slow on-rate or because slowly growing microtubule ends have a different, possibly more pre-catastrophe-like structure. Thus, CSPP-L stabilizes microtubule polymerization intermediates at microtubule tips or damage sites when tubulin addition occurs slowly.

**Figure 6:**
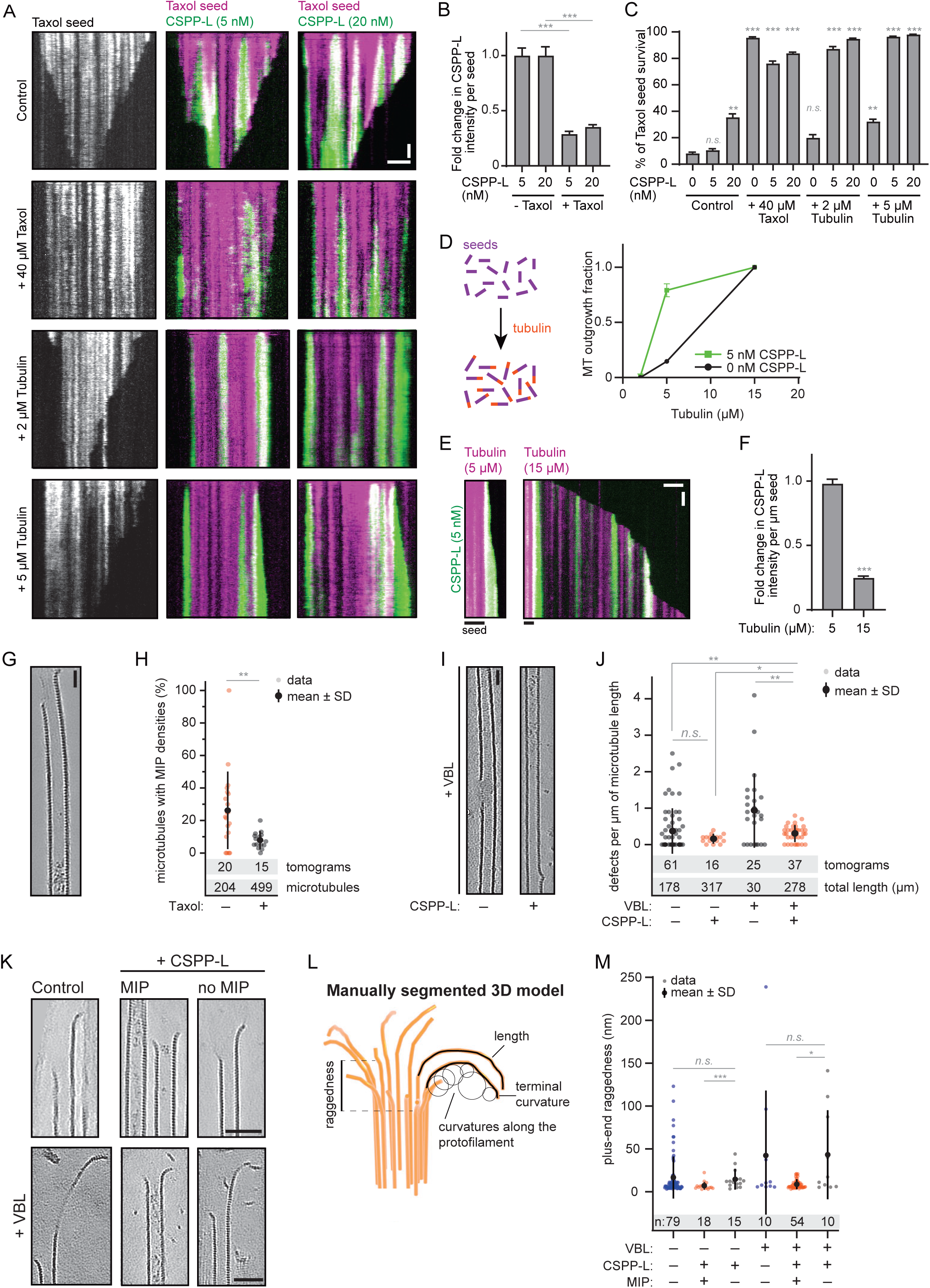
**CSPP1 stabilizes microtubules by promoting lattice repair** A. Kymographs of Taxol-stabilized microtubule seeds in absence or presence of the indicated Taxol, tubulin and GFP-CSPP-L concentrations. Scale bars, 2 μm (horizontal) and 60 s (vertical). B. GFP-CSPP-L intensity quantification per microtubule seed (from kymographs as shown in A). Mean GFP-CSPP-L intensity was measured along the complete seed 2 min after flowing in the protein. The average mean intensity of GFP-CSPP-L in presence of 40 µM Taxol was normalized to the average mean intensity in absence of free Taxol. Number of Taxol-stabilized microtubule seeds analyzed: 5 nM CSPP-L alone, n=102; 20 nM CSPP-L alone, n=108; 40 µM Taxol, n=99; 40 µM Taxol together with 5 nM CSPP-L, n=84; 40 µM Taxol together with 20 nM CSPP-L, n=114. Error bars represent SEM. ***, p< 0.001; n.s., not significant; Kruskal-Wallis test followed by Dunńs post- test. Data quantified from two experiments. C. Quantification of the percentage of microtubule seeds which survived 5 min after flow-in of the reaction mix (from kymographs as shown in A). Number of Taxol-stabilized microtubule seeds analyzed: control: n=95; 5 nM CSPP-L alone, n=120; 20 nM CSPP-L alone, n=110; 40 µM Taxol, n=99; 40 µM Taxol together with 5 nM CSPP-L, n=84; 40 µM Taxol together with 20 nM CSPP-L, n=120; 2 µM tubulin alone, n=115; 2 µM tubulin together with 5 nM CSPP-L, n=124; 2 µM tubulin together with 20 nM CSPP-L, n=112; 5 µM tubulin alone, n=122; 5 µM tubulin together with 5 nM CSPP-L, n=106; 5 µM tubulin together with 20 nM CSPP-L, n=128. Error bars represent SEM. ***, p< 0.001; **, p< 0.01; n.s., not significant; Kruskal-Wallis test followed by Dunńs post-test. In “control”, conditions with 5 and 20 nM CSPP-L are compared to 0 nM CSPP-L, and for all other bars, comparisons were made to the same CSPP-L concentration in the control condition. Data quantified from two experiments. D. Quantification of the fraction of the total GMPCPP seeds that showed microtubule outgrowth within 10 min at indicated tubulin concentrations, with tubulin alone or together with 5 nM GFP- CSPP-L. Number of GMPCPP seeds analyzed: 2 µM tubulin alone, n=74; 5 µM tubulin alone, n=75; 15 µM alone, n=69; 2 µM tubulin together with 5 nM CSPP-L, n=70; 5 µM tubulin together with 5 nM CSPP-L, n=66; 15 µM tubulin together with 5 nM CSPP-L FL, n=71. Error bars represent SEM. Data quantified from two experiments. E. Kymographs of GMPCPP-stabilized microtubule seeds in the presence of 5 nM GFP-CSPP-L and the indicated tubulin concentrations. Scale bars, 2 μm (horizontal) and 60 s (vertical). F. GFP-CSPP-L intensity quantification per µm newly grown microtubule lattice (from kymographs as shown in E). GFP-CSPP-L integrated intensity was measured on newly grown lattice 5 min after flow-in of the reaction mix. Integrated intensity was normalized to newly grown microtubule lattice length, and the average mean intensity of GFP-CSPP-L in presence of 15 µM tubulin was normalized to the average mean intensity in presence of 5 µM tubulin. Number of growth episodes analyzed: 5 µM tubulin, n=105; 15 µM tubulin, n=104. Error bars represent SEM. ***, p< 0.001, Mann-Whitney test. Data quantified from two experiments. G. Denoised tomograms of dynamic microtubules polymerized in the presence of 250 µM Taxol, resuspended in buffer containing only 20 nM GFP-CSPP-L with or without free 40 µM Taxol, vitrified on EM grids. Scale bar, 25 nm. H. Quantification of the percentage of microtubules containing luminal densities from total microtubules (from tomograms as shown in G). Orange and grey dots (single data points, tomograms), black circle (mean), SD (error bars). **, p< 0.01, Mann-Whitney test. Analysis from two experiments. I. Denoised tomograms of dynamic microtubules polymerized in the presence or absence of 250 nM Vinblastin with or without 20 nM GFP-CSPP-L, vitrified on EM grids. Scale bar, 25 nm. J. Quantification of the number of defects per µm microtubule (from tomograms as shown in I). Orange and grey dots (single data points, tomograms), black circles (mean), SD (error bars). *, p<0.1, **, p< 0.01, n.s., not significant, Mann-Whitney test. Analysis from two experiments. K. Denoised tomograms of microtubule ends in presence or absence of 250 nM Vinblastin with or without 20 nM GFP-CSPP-L, vitrified on EM grids. Scale bars, 50 nm. L. Parameters extracted from manual segmentations of terminal protofilaments. M. Quantification of plus-end raggedness (from tomograms as shown in K). Blue, orange and grey dots (single data points, tomograms), black circle (mean), SD (error bars). *, p<0.1, ***, p< 0.001, n.s., not significant, Mann-Whitney test. Analysis from two experiments. See also Figure S4.

To better understand the mechanism underlying the activity of CSPP1, we again turned to cryo-ET. We stabilized microtubules by the addition of Taxol, and then resuspended them in a buffer containing CSPP-L with or without free Taxol, and no free tubulin. Absence of free Taxol increased CSPP-L binding: on average 26% ± 23% of microtubules contained MIP densities, compared to only 8% ± 5% in presence of Taxol (p < 0.01, Fig. 6G, H). Despite the fact that samples with disassembling Taxol-stabilized microtubules contained many incomplete lattices and tubulin sheets, we only observed MIP densities inside fully closed tubes (Fig. 6G). Moreover, CSPP-L accumulation zones did not bind CAMSAP3 in in vitro assays and thus did not contain lattice apertures (Fig. S4C), unlike previous observations with Fchitax-3 (Rai et al., 2020). Presence of vinblastine during microtubule growth led to the presence of more numerous defects in the microtubule lattices (Fig. 6I, J). However, presence of both vinblastine and CSPP-L during microtubule growth led to a significant reduction of the number of lattice defects, comparing to vinblastine alone (0.3 ± 0.2 vs 1 ± 1 defects/µm, *p =* 0.005). These observations, in combination with increased number of MIP-containing microtubules in presence of vinblastine (Fig. 4A, B), support our hypothesis that CSPP1 can enter microtubules through lattice openings, and then promote their repair.

In order to explain how CSPP1 stabilizes microtubules, we analyzed the shapes of terminal tubulin flares in our cryo-ET samples. We used manual segmentation to extract parameters of protofilament shape and length in 3D, as well as raggedness, or tapering (Fig. 6K, L). Comparing microtubule ends with MIPs to MIP-free microtubules in the same sample, we did not observe any significant differences in protofilament length or curvature (Fig. S4D). However, we did observe reduced raggedness of MIP-containing microtubule ends comparing MIP-positive and MIP-negative microtubules in presence of both CSPP-L and vinblastine (Fig. 6M). This might indicate CSPP1 does no act at terminal protofilament flares, but stabilizes microtubules by holding protofilaments together within the tube, thus preventing microtubule disassembly and allowing them to resume growth. In a similar way, CSPP1 could potentially bind to damaged lattices and hold protofilaments together, to enable lattice repair by tubulin incorporation.

## Discussion

While a lot of information exists about the control of microtubule dynamics by proteins associated with the outer microtubule surface, the regulatory effects of factors binding to microtubule lumen are understood much less well. Here, we show that the ciliary tip regulator CSPP1 is a MIP and dissect its behavior and molecular function. We show that CSPP1 displays some striking parallels to microtubule-stabilizing compounds, such as taxanes and epothilones, which also bind to microtubule lumen (reviewed in (Steinmetz and Prota, 2018)). Similar to these compounds, CSPP1 binds to polymerizing microtubule ends in the pre-catastrophe state, when the GTP cap is diminished, prevents catastrophe and induces microtubule pausing followed by growth; at low concentration, CSPP1 triggers formation of sites of stabilized microtubule lattice that cause repeated rescues (“stable rescue sites” (Rai et al., 2020)). Preferential accumulation of CSPP1 at growing microtubule ends can be explained by the better accessibility of intraluminal binding sites, which become available when tubulin dimers are added to microtubule ends. Theory predicts that intraluminal diffusion of a protein with affinity for the inner microtubule surface would be very slow (Odde, 1998). Furthermore, unlike small molecules, CSPP1 would be too large to penetrate into the microtubule lumen through the regular lattice fenestrations, although it does bind to sites where the lattice has been damaged. Additionally, for CSPP1 to be able to accumulate inside the microtubule, this damage needs to be sufficiently large as accumulations are only observed in Taxol-stabilized microtubules with large defects but not in GMPCPP-stabilized microtubules which have smaller defects. The selectivity of CSPP1 for pre-catastrophe microtubule ends could be explained by their specific conformation (such as presence of tubulin sheets or tapers, or the loss of GTP-tubulin) or simply by their slow growth. The observation that CSPP1 binds to growing microtubule ends better when tubulin concentration is low and that it strongly accumulates inside microtubules lattices polymerized from the minus-end is in agreement with its preference for slowly polymerizing microtubule ends. The striking overlap between the binding profiles of CSPP1 and the fluorescent taxane Fchitax-3, combined with our previous data demonstrating cooperative Fchitax-3 binding to microtubule ends (Rai et al., 2020), suggests that also CSPP1 can cooperatively bind to microtubule tips that undergo a growth perturbation. This would explain how CSPP1 forms regions of high enrichment when present at low concentrations. After binding, CSPP1 exerts a microtubule-stabilizing effect by preventing shrinkage; it could do so by supporting individual protofilaments and/or by promoting lateral interactions between protofilaments, and both mechanisms would be consistent with the action of microtubule-stabilizing agents ((Elie-Caille et al., 2007; Prota et al., 2013), reviewed in (Steinmetz and Prota, 2018)). Spanning lateral protofilament contacts could potentially explain how CSPP1 reduces tip raggedness and why it is not found at protofilament flares.

Another interesting property of CSPP1 is its ability to induce pausing. While this property also resembles the effect of low concentrations of taxanes, in CSPP1, the lumen binding and growth-inhibiting functions depend on two separate protein domains. Presence of two activities, an activity that inhibits polymerization and an activity that prevents microtubule shrinkage, seems to be a common property of microtubule growth inhibitors, such as the kinesin-4 KIF21B (van Riel et al., 2017) or the centriolar protein CPAP (Sharma et al., 2016). In CSPP1, both regulatory domains are predicted to be helical and are quite short, less than a hundred amino acids. Presence of α-helices seems to be a common property of ciliary MIPs, including many linearly arranged proteins that form the regularly spaced inner sheath within ciliary doublets (Gui et al., 2021; Ichikawa and Bui, 2018; Ma et al., 2019). Identification of a minimal lumen-binding domain of CSPP1 (termed here the MTB) can be potentially useful for directing different protein activities to the microtubule lumen. It is possible that the binding site of the CSPP1 MTB domain overlaps with that of Taxol, because we found some evidence of competition between Taxol and CSPP1 in microtubule stabilization assays. Importantly, there is also a notable difference between the effects microtubule-stabilizing drugs and CSPP1: taxanes induce structural defects (holes) in microtubule lattices because they promote switching in protofilament number (Rai et al., 2021). In contrast, CSPP1 seems to promote lattice integrity. Although CSPP1 can specifically bind to the sites of lattice damage, CSPP1 densities are predominantly found within complete tubes; moreover, CSPP1 reduces the number of vinblastine-induced lattice defects and stabilizes eroding microtubule seeds. CSPP1 likely acts in part by stabilizing protofilament ends close to the damage sites and possibly by promoting tubulin incorporation to form complete tubes. Whether CSPP1 participates in repair of microtubule defects in cells, either on cytoplasmic or axonemal microtubules, remains to be determined. There are indications that cellular microtubules can be damaged by interaction with other microtubules, severing enzymes or motor proteins that use microtubules as rails (Aumeier et al., 2016; Gazzola et al., 2022; Triclin et al., 2021; Vemu et al., 2018). The ability of CSPP1 to specifically bind to incomplete microtubules can be harnessed for studying microtubule damage and repair. It should be noted that another protein, SSNA1, was also reported to bind to microtubule defects, albeit it appears much less potent than CSPP1 in stabilizing microtubules, because 0.5-5 µM SSNA1 was needed to affect microtubule growth in vitro (Lawrence et al., 2021), whereas CSPP1 displays strong effects already at 5-10 nM concentration. It would be interesting to examine whether SSNA1 is also a MIP, as it was reported to stabilize partial microtubule structures (Basnet et al., 2018).

CSPP1 also shows some similarities to another intraluminal protein that has been analyzed in vitro, MAP6 (Cuveillier et al., 2020). While MAP6 shows some strikingly distinct features, such as the induction of microtubule coiling and lattice apertures (Cuveillier et al., 2020), both MAP6 and CSPP1 are microtubule stabilizers, which reduce overall microtubule shrinkage and promote rescues. Furthermore, both proteins contain a short domain that can perturb processive growth. In the case of MAP6, this domain is also required for the formation of intraluminal particles, and without it, the protein seems to function on the outer microtubule surface. In contrast, our Cryo-ET and MINFLUX data support the idea that CSPP1 binds only to the inner surface of the microtubule. It is however still possible that some parts of CSPP1 extend out of the tube. For example, the site of action of the growth-inhibiting part of CSPP1 is currently unclear, as the shape and curvature of the protofilament flares in the presence of CSPP1 looked very similar to that of control microtubules and thus provided no clues on the nature of this activity. Furthermore, CSPP1 is part of a multiprotein module associated with ciliary tips (Latour et al., 2020), and two other members of the same module, TOGARAM1 and CEP104, are likely to bind to the outer microtubule surface, because they contain canonical tubulin-binding TOG domains; moreover, CEP104 binds to EBs, which decorate microtubules from the outside (Al-Jassar et al., 2017; Das et al., 2015; Jiang et al., 2012; Rezabkova et al., 2016).

CSPP1 participates in controlling the elongation and stability of ciliary axonemes, and when CSPP1 or its binding partners are absent, ciliogenesis is impaired and cilia are shorter (Frikstad et al., 2019; Latour et al., 2020; Patzke et al., 2010). Our findings help to explain the microtubule-stabilizing activity of CSPP1 and suggest that ciliary tips are kept in shape by protein complexes that span both the inner and the outer microtubule surface. This arrangement might be important for controlling different signaling pathways such as Hedgehog signaling, which strongly relies on the state of axoneme tip and is dysregulated by ciliopathies (Andreu-Cervera et al., 2021; Hildebrandt et al., 2011; Reiter and Leroux, 2017). Furthermore, the similarity between the activities of CSPP1 and microtubule-stabilizing agents raise an interesting possibility that the absence of CSPP1 or its binding partners might be compensated by such compounds, suggesting potential avenues for pharmacological intervention in ciliopathies.

## Author Contributions

C.M. vdB designed and performed protein purifications and in vitro reconstitution experiments, analyzed data and wrote the paper; V.A.V. designed and performed cryo-ET experiments, analyzed data and wrote the paper, S.S., Z.H. and T.Z facilitated and performed MINFLUX imaging and analysis, K.E.S. performed and analyzed mass spectrometry experiments, I.G. performed and analyzed live cell imaging experiments using TIRF microscopy, S.P. provided reagents, insights into data analysis and contributed to writing, M.D. coordinated the project, A.A. coordinated the project and wrote the paper.

## Acknowledgements

We thank A. Jakobi (TU Delft) for providing access to a GPU computational cluster. V.A.V. acknowledges support from the QMUL Startup grant (SBC8VOL2). This work was supported by the European Research Council Synergy grant 609822 to M.D. and A.A. and the ZonMW TOP 91216006 program. We acknowledge the access and services provided by the Imaging Centre at the European Molecular Biology Laboratory (EMBL IC), generously supported by the Boehringer Ingelheim Foundation. The MINFLUX image acquisition of in vitro microtubules was facilitated by the Christian Boulin Fellowship awarded to C.M. vdB.

## Methods

### DNA constructs, cell lines and cell culture

CSPP1 truncations expressed in mammalian cells were made from the full-length constructs described previously (Patzke et al., 2005; Patzke et al., 2006) in modified pEGFP-C1 or pmCherry- C1 vectors with a StrepII tag. HEK293T cells and COS-7 cells (ATCC) were cultured in DMEM medium (Lonza) supplemented with 10% fetal calf serum (FCS) (GE Healthcare Life Sciences) and 1% (v/v) penicillin/streptomycin. All cells were routinely checked for mycoplasma contamination using the MycoAlertTM Mycoplasma Detection Kit (Lonza). For overexpression of CSPP1 constructs, COS-7 cells were transiently transfected with FuGENE6 (Promega) with different StrepII-GFP-CSPP1 constructs for 24 hours. Single transfections were used for immunofluorescence experiments and co-transfections with EB3-mCherry (Stepanova et al., 2003), βIVb-tubulin-mCherry (Bouchet et al., 2016) or StrepII-GFP-MAP7 FL (Hooikaas et al., 2019) were used for live-cell imaging or MINFLUX microscopy.

### Protein purification from HEK293T cells for in vitro reconstitution assays

For the purification of CSPP1 constructs, HEK293T cells were transiently transfected with polyethyleneimine (Polysciences) with different StrepII-GFP-CSPP1 constructs. The cells were harvested 28 hours after transfection. Cells from a 15 cm dish were lysed in 500 µl lysis buffer (50 mM HEPES, 300 mM NaCl, 1 mM MgCl_2_, 1 mM DTT, 0.5% Triton X-100, pH 7.4) supplemented with protease inhibitors (Roche) on ice for 15 minutes. The lysate was cleared from debris by centrifugation and the supernatant was incubated with 20 µl StrepTactin beads (GE Healthcare) for 45 min. Beads were washed five times with a 300 mM salt wash buffer (50 mM HEPES, 300 mM NaCl, 1 mM MgCl_2_, 1 mM EGTA, 1 mM DTT, 0.05% Triton X-100, pH 7.4) and three times with a 150 mM salt wash buffer (similar to the 300 mM salt buffer but with 150 mM NaCl). The protein was eluted in elution buffer (similar to the 150 mM salt wash but supplemented with 2.5 mM d-Desthiobiotin (Sigma-Aldrich)) where the volume depended on the expression levels before harvesting. Purified proteins were snap-frozen and stored at -80°C.

### Mass spectrometry

To confirm we purified GFP-CSPP-L without any interacters that could affect its effect on microtubule dynamics, the purified protein sample was digested using S-TRAP microfilters (ProtiFi) according to the manufacturer’s protocol. In short, 7 µg of protein sample was denatured in 5% SDS buffer and reduced and alkylated using DTT (20 mM, 10 min, 95°C) and iodoacetamide (IAA; 40 mM, 30 min). After acidification, the proteins were precipitated using a methanol triethylammonium bicarbonate buffer (TEAB) after which they were loaded on the S-TRAP column. The trapped proteins were washed four times with the methanol TEAB buffer and then digested using 1 µg Trypsin (Promega) overnight at 37°C. Digested peptides were eluted and dried in a vacuum centrifuge before liquid chromatography-mass spectrometry (LC-MS) analysis.

The sample was analyzed by reversed-phase nLC-MS/MS using an Ultimate 3000 UHPLC coupled to an Orbitrap Q Exactive HF-X mass spectrometer (Thermo Scientific). Digested peptides were separated using a 50 cm reversed-phase column packed in-house (Agilent Poroshell EC-C18, 2.7 µm, 50cm x 75 µm). The peptides were eluted from the column at a flow rate of 300 nl/min using a linear gradient with buffer A (0.1% formic acid (FA)) and buffer B (80% acetonitrile (ACN), 0.1% FA) ranging from 13-44% B over 38 min. This procedure was followed by a column wash and re-equilibration step resulting in a ttotal data acquisition time of 55 min. Mass spectromtery data were acquired using a data-dependent aqcuisition (DDA) method with the following MS1 scan parameters: maximum injection time of 20 msec, automatic gain control (AGC) target equal to 3E6, 60,000 resolution, the scan range of 375-1600 m/z, acquired in profile mode. The MS2 method was set at 15,000 resolution, an automatic maximum injection time, with an AGC target set to standardand an isolation window of 1.4 m/z. Scans were acquired using a fixed first mass of 120 m/z and a mass range of 200-2000, and an normalized collision energy (NCE) of 28. Precursor ions were selected for fragmentation using a 1-second scan cycle, a dynamic exclusion time set to 10 sec, and a precursor charge selection filter for ions possessing +2 to +6 charges.

Raw files were processed using Proteome Discoverer (PD) (version 2.4, Thermo Scientific). MSMS fragment spectra were searched using Sequest HT against a human database (UniProt, year 2020) that was modified to contain the exact protein sequence from SII-GFP-CSPP-L and a common contaminants database. The search parameters were set using a fragment mass tolerance of 0.06 Da and a precursor mass tolerance of 20 ppm and. The maximum amount of missed cleavages for trypsin digestion was set to two. Methionine oxidation and protein N-term acytelyation were set as variable modifications and carbamidomethylation was set as a fixed modification. Percolator was used to assign a 1% false discovery rate (FDR) for peptide spectral matches, and a 1% FDR was applied to protein and peptide assemblies. For peptide-spectrum match (PSM) inlcusion, an additional filter was set to require a minimum Sequest score of 2.0. The Precursor Ion Quantifier node was used for MS1 based quantification, default were settings applied. Precursor ion feature matching was enabled using the Feature Mapper node. Proteins that matched the common contaminate database were filtered out from the results table.

### In vitro reconstitution assays

#### Microtubule seed preparation

Double-cycled GMPCPP-stabilized microtubule seeds or Taxol-stabilized microtubule seeds were used as templates for microtubule nucleation or to test protein binding in in vitro assays. GMPCPP-stabilized microtubule seeds were prepared as described before (Mohan et al., 2013). Briefly, a tubulin mix consisting of 70% unlabeled porcine brain tubulin, 18% biotin-labeled porcine tubulin and 12% rhodamine-labeled porcine tubulin (all from Cytoskeleton) was incubated with 1 mM GMPCPP (Jena Biosciences) at 37°C for 30 minutes. Polymerized microtubules were pelleted by centrifugation in an Airfuge for 5 min at 199,000 x g and then depolymerized on ice for 20 min. Next, microtubules were let to polymerize again at 37°C with newly added 1 mM GMPCPP. Polymerized microtubule seeds were then pelleted as above and diluted tenfold in MRB80 buffer containing 10% glycerol. Last, microtubule seeds were frozen and stored at -80°C. Taxol-stabilized microtubule seeds were prepared as described before with some modifications (Aher et al., 2020). Briefly, a tubulin mix consisting of 28 µM porcine brain tubulin, 10% biotin-labeled porcine tubulin and 4.5% rhodamine-labeled porcine tubulin was incubated with 2 mM GTP (Sigma-Aldrich) and 20 µm Taxol at 37°C for 35 minutes. Then, 20 µM Taxol was added to the tubulin mix and polymerized microtubules were pelleted by centrifugation for 15 min at 16,200 x g at room temperature. The microtubule pellet was resuspended in warm 20 µM Taxol solution in MRB80 buffer and stored at room temperature in the dark for a maximum of one day.

#### In vitro reconstitution assays

In vitro assays with dynamic or stabilized microtubules were performed as described before (Rai et al., 2020). In short, plasma-cleaned glass coverslips (square or rectangular) were attached on microscopic slides by two strips of double-sided tape. The coverslips were functionalized by sequential incubation with 0.2 mg/ml PLL-PEG-biotin (Susos AG, Switzerland) and 1 mg/ml neutravidin (Invitrogen) in MRB80 buffer (80 mM piperazine-N, N[prime]-bis (2-ethane sulfonic acid), pH 6.8, supplemented with 4 mM MgCl_2_, and 1 mM EGTA). Then, GMPCPP- or Taxol-stabilized microtubule seeds were attached to the coverslips through biotin-neutravidin interactions. During the subsequent blocking step with 1 mg/ml κ-casein, the reaction mix containing the different concentrations of purified proteins and drugs was spun down in an Airfuge for 5 min at 119,000 x g. For dynamic microtubules, the reaction mix consisted of MRB80 buffer supplemented with 15 µM porcine brain tubulin (100% dark porcine brain tubulin when 20 nM GFP-EB3 or mCherry-EB3 was added, or 97% dark porcine brain tubulin with 3% rhodamine- or HiLyte488-labeled porcine tubulin), 50 mM KCl, 1 mM GTP, 0.2 mg/ml κ-casein, 0.1% methylcellulose and oxygen scavenger mix [50 mM glucose, 400 µg ml−1 glucose oxidase, 200 µg/ml catalase and 4 mM DTT]. For stabilized microtubules, porcine tubulin, GTP and EB3 were omitted from the reaction mix. After spinning, the reaction mix was added to the flow chamber and the flow chamber was most often sealed with vacuum grease or left open (for flow-in assays during acquisition or for MINFLUX sample preparation). Microtubules were imaged immediately at 30°C using a total internal reflection fluorescence (TIRF) microscope. All tubulin products were from Cytoskeleton Inc.

To estimate the number of GFP-CSPP-L molecules per 8 nm microtubule, two parallel flow chambers were made on the same coverslip. In one chamber, regular microtubule dynamic assay in the presence of GMPCPP-stabilized microtubule seeds with tubulin, EB3-mCherry and 5 nM GFP-CSPP-L was performed. The other chamber was incubated with strongly diluted GFP protein so that single molecules were detectable. Microtubules were let to polymerize for 5-10 minutes. Then, for both the chamber with single GFP molecules and the chamber with dynamic microtubule and GFP-CSPP-L, 20 images of unexposed coverslip areas were acquired at 100-ms exposure time using high laser intensity.

#### In vitro assays for Cryo-ET sample preparation

Sample preparation for imaging in vitro microtubules with cryo-ET is a slightly modified version of the method described above. All steps occur in a tube instead of a flow chamber. After centrifugation of the reaction mix for dynamic microtubules, GMPCPP-stabilized seeds and 5 nm gold particles were added, and microtubules were let to polymerize for 20-30 min at 37°C. Then, 3.5 µl was transferred to a recently glow-discharged, lacey carbon grid suspended in the chamber of Leica EM GP2 plunge freezer, equilibrated at 37°C and 98% relative humidity. The grid was immediately blotted for 4 s and plunge-frozen in liquid ethane.

#### In vitro assays for MINFLUX sample preparation

Sample preparation for imaging in vitro microtubules with MINFLUX microscopy is a slightly modified version of the method described above. For flow-chambers, round plasma-cleaned coverslips were attached to big, rectangular coverslips via two stripes of glue (Twinsil®). The reaction mix contained the same components as for dynamic microtubules, supplemented with an CF680-GFP-Nanobody and SNAP-Abberior FLUX-640. After addition of the reaction mix, the chamber was left open and was incubated in a 30°C incubator for 15 min. To remove background signal, the flow chamber was washed with a second reaction mix containing 25 µM tubulin before fixing with 1% glutaraldehyde (Electron Microscopy Sciences) for 5 minutes at room temperature. After washing with MRB80, the round glass coverslip was demounted and stored in MRB80 at 4°C or incubated with gold nanoparticles (Nanopartz) for 5 min. Then, the coverslips were mounted in GLOX buffer (50 mM Tris/HCl pH 8, 10 mM NaCl, 10% (w/v) d-glucose, 500 µg/ml glucose oxidase, 40 µg/ ml glucose catalase) supplemented with 56 mM 2-Mercaptoethylamin (MEA) and sealed with glue (Picodent Twinsil®).

### Immunofluorescence staining of fixed cells

#### Sample preparation for widefield fluorescence imaging

For immunofluorescence staining experiments, COS-7 cells were seeded on coverslips one day before transfection. Cells were fixed after 24 hours with either −20°C MeOH for 10 min (staining for acetylated tubulin, α-tubulin, PCM1 and CSPP1) or −20°C MeOH for 10 min followed by 4% paraformaldehyde for 15 min at room temperature (staining for α-tubulin and EB1). This was followed by permeabilization with 0.15% Triton X-100 for 2 min. Next, samples were blocked with 1% bovine serum albumin (BSA) diluted in phosphate buffered saline (PBS) supplemented with 0.05% Tween-20 for 45 min at room temperature and sequentially incubated with primary antibodies for 1 hour at room temperature and fluorescently labeled with secondary antibodies for 45 min at room temperature. Finally, samples were washed, dried and mounted in Vectashield (Vector laboratories).

#### Sample preparation for MINFLUX microscopy imaging

For immunofluorescence staining experiments, COS-7 cells were seeded on coverslips one day before transfection. 24 hours after transfection, the cells were incubated with warm extraction buffer (0.2% glutaraldehyde, 0.35% Triton X-100 in MRB80) for 2 min before incubation with fixation buffer (0.1% glutaraldehyde, 4% paraformaldehyde and 4% sucrose (w/v)) for 10 min at room temperature (staining for α-tubulin and EB1) followed by permeabilization with 0.5% Triton X-100 for 10 min. Next, samples were quenched with 100 mM NaBH_4_ before blocking with Image-iT Signal Enhancer (Thermo Fisher Scientific) for 30 min at room temperature and sequentially incubated with 1 µM Alexa647-SNAP-dye (NEB) and 1 mM DTT in PBS for 1 hour at room temperature. Next, samples were blocked with 1% bovine serum albumin (BSA) diluted in phosphate buffered saline (PBS) for 50 min at room temperature and sequentially incubated with CF680-GFP-Nanobody for 1 hour at room temperature or overnight at 4°C. Coverslips were incubated with gold nanoparticles (Nanopartz) for 5 min. Then, the coverslips were mounted in GLOX buffer supplemented with 56 mM MEA and sealed with glue (Picodent Twinsil®).

### Microscopy

#### Widefield microscopy

Fixed and stained COS-7 cells were imaged using widefield fluorescence illumination on a Nikon Eclipse Ni upright microscope equipped with a Nikon DS-Qi2 camera (Nikon), an Intensilight C-HGFI precentered fiber illuminator (Nikon), ET-DAPI, ET-EGFP and ET-mCherry filters (Chroma), controlled by Nikon NIS Br software and using a Plan Apo Lambda 60x NA 1.4 oil objective (Nikon). For presentation, images were adjusted for brightness using ImageJ 1.50b.

#### TIRF microscopy

In vitro reconstitution assays and live COS-7 cells overexpressing GFP-CSPP-L and mCherry-EB3 were imaged on previously described (iLas2) TIRF microscope setups (Aher et al., 2020).

In brief, we used an inverted research microscope Nikon Eclipse Ti-E (Nikon) with the perfect focus system (Nikon), equipped with Nikon CFI Apo TIRF 100x 1.49 N.A. oil objective (Nikon) and controlled with MetaMorph 7.10.2.240 software (Molecular Devices). The microscope was equipped with TIRF-E motorized TIRF illuminator modified by Gataca Systems (France). To keep the in vitro samples at 30°C, a stage top incubator model INUBG2E-ZILCS (Tokai Hit) was used. For excitation, 490 nm 150 mW Vortran Stradus 488 laser (Vortran) and 561 nm 100 mW Cobolt Jive (Cobolt) lasers were used. We used ET-GFP 49002 filter set (Chroma) for imaging of proteins tagged with GFP or tubulin labeled with Hylite488 or ET-mCherry 49008 filter set (Chroma) for imaging of proteins tagged with mCherry or tubulin labeled with rhodamine. Fluorescence was detected using an Prime BSI camera ( Teledyne Photometrics) with the intermediate lens 2.5X (Nikon C mount adaptor 2.5X) or an EMCCD Evolve 512 camera (Roper Scientific) without an additional lens. The final resolution using Prime BSI camera was 0.068 μm/pixel, using EMCCD camera it was 0.063 μm/pixel.

The iLas3 system (Gataca Systems (France)) is a dual laser illuminator for azimuthal spinning TIRF (or Hilo) illumination and targeted photomanipulation option. This system was installed on Nikon Ti microscope (with the perfect focus system, Nikon), equipped with 489 nm 150 mW Vortran Stradus 488 laser (Vortran) and 100 mW 561 nm OBIS laser (Coherent), 49002 and 49008 Chroma filter sets, EMCCD Evolve DELTA 512 camera (Teledyne Photometrics) with the intermediate lens 2.5X (Nikon C mount adaptor 2.5X), CCD camera CoolSNAP MYO (Teledyne Photometrics) and controlled with MetaMorph 7.10.2.240 software (Molecular Device). To keep the in vitro samples at 30°C or the live cells at 37°C, a stage top incubator model INUBG2E-ZILCS (Tokai Hit) was used. The final resolution using EMCCD camera was 0.064 μm/pixel, using CCD camera it was 0.045 μm/pixel. This microscope was also used for photoablation. The 532 nm Q-switched pulsed laser (Teem Photonics) as part of iLas3 system was used for photoablation by targeting the laser on the TIRF microscope very close but not directly at the microtubule lattice to induce damage. For photodamage, a circle with a diameter of 7 pixels was used for 50 ms illumination at 20%–25% laser power of the 532 nm pulsed laser.

#### Spinning disk microscopy

Photodamage assays in cells were performed using spinning disk microscopy. COS-7 cells overexpressing GFP-CSPP-L and β-tubulin-mCherry were imaged using confocal spinning disc fluorescence microscopy on an inverted research microscope Nikon Eclipse Ti-E (Nikon), equipped with the perfect focus system (Nikon), Nikon Plan Apo VC 100x N.A. 1.40 oil objective (Nikon) and a spinning disk-based confocal scanner unit (CSU-X1-A1, Yokogawa). The system was also equipped with ASI motorized stage with the piezo plate MS-2000-XYZ (ASI), Photometrics PRIME BSI sCMOS camera (Teledyne Photometrics) and controlled by the MetaMorph 7.10.2.240 software (Molecular Devices). For imaging we used 487 nm 150 mW Vortran Stradus 488 (Vortran) and 100 mW 561 nm OBIS (Coherent) lasers, the ET-EGFP/mCherry filter (Chroma) for spinning-disc-based confocal imaging. The final resolution using PRIME BSI camera was 0.063 μm/pixel. To keep the live cells at 37°C, a stage top incubator model INUBG2E-ZILCS (Tokai Hit) was used. The 355 nm laser (Teem Photonics) of the iLAS pulse system was used to induce photodamage by targeting the laser on the spinning disk microscope in a 1 pixel thick line across microtubules in the z-plane under the nucleus at 9%-11% laser power to induce damage.

#### MINFLUX microscopy

MINFLUX imaging was performed on an Abberior MINFLUX microscope (Abberior) equipped with a 1.4 NA 100× Oil objective lens as previously described (Schmidt et al., 2021). Two color images were recorded using ratiometric detection on two avalanche photodiodes. The fluorescence signal of two far-red fluorophores was split at 685 nm into two detection channels (Ch1: 650-685 nm and Ch2: 685-750 nm), the ratio between both detector channels allowed to assign the individual single molecule events to the respective fluorophores. Images were acquired in 2D or 3D MINFLUX imaging mode using a 642 nm excitation laser (17.4 μW cm–2). Laser powers were measured at the position of the objective back focal plane using a Thorlabs PM100D power meter equipped with a S120C sensor head.

#### Cryo-ET microscopy

Images were recorded on a JEM3200FSC microscope (JEOL) equipped with an in-column energy filter operated in zero-loss imaging mode with a 30 eV slit. Movies consisting of 8-10 frames were recorded using a K2 Summit direct electron detector (Gatan), with a target total electron dose of 80 e−/Å2. Images were recorded at 300 kV with a nominal magnification of 10000, resulting in a pixel size of 3.668 Å at the specimen level. Imaging was performed using SerialEM software (Mastronarde, 2005), recording bidirectional tilt series starting from 0° ±60°; tilt increment 2°; target defocus -4 µm.

### Image analysis

#### Analysis of microtubule plus end dynamics in vitro

Movies of dynamic microtubules, acquired as describe above, were corrected for drift, and kymograps were generated using the ImageJ plugin KymoResliceWide v.0.4 (https://github.com/ekatrukha/KymoResliceWide). The microtubule tips was traced with lines and measured lengths and angles were used to calculate the microtubule dynamics parameters such a sgrowth rate, pause duration, event duration and all transition events. All events with growth rates faster than 0.24 µm/min were categorized as growth events and all events with shrinkage rates faster than 0.24 µm/min were categorized as shrinkage events. The events with slower growth rates or faster shrinkage rates than the before mentioned rates were categorized as pause events. Only growth events longer than 0.40 µm and pause events longer than 20 seconds were included in the analysis. Transition frequency was calculated by dividing the sum of the transition events per experiment by the total time this event could have occurred.

#### Quantification of EB1 comets

Images of COS-7 cells overexpressing GFP-CSPP1 constructs and stained for α-tubulin and EB1 were acquired on an widefield microscope as described above. The background was subtracted using the Rollin Ball Background Substraction plugin in ImageJ. This plugin uses the rolling-ball algorithm where we set the radius to 10 pixels. EB1 comets were detected by “MaxEntropy” thresholding and subsequent particle analysis with a minimal size cut-off of 0.10 µm^2^ and the total number of EB1 comets per cell was normalized to 100 µm^2^.

#### 3D volume reconstruction and analysis

Reconstruction, denoising, and analysis of tomographic volumes were performed as described previously (Ogunmolu et al., 2021). In brief, direct electron detector movie frames were aligned using MotionCor2 (Zheng et al., 2017) and then split into even and odd stacks used further for denoising. Tilt series alignment and tomographic reconstructions were performed with IMOD 4.11 (Kremer et al., 1996). Final tomographic volumes were binned by two, corrected for contrast transfer function, and the densities of gold beads were erased in IMOD. CryoCARE denoising was performed on tomograms reconstructed from the same tilt-series using even and odd movie frames (Buchholz et al., 2019). Tubulin lattice defects were identified upon visual inspection of denoised tomograms in 3dmod as interruptions of regular microtubule lattice that could not attributed to missing wedge artifacts. Microtubules were sometimes damaged at microtubule-carbon or microtubule-microtubule contacts followed by blotting; these instances were not included in the quantification of defects.

Automated segmentation of denoised tomograms into tubulin and MIP densities was performed using the tomoseg module of EMAN2.2 (Chen et al., 2017). To do this, we trained three separate neural networks: ‘microtubules’, ‘tubulin’ and ‘MIP’. The resulting segmentations were used to mask the denoised tomographic densities using UCSF Chimera (Pettersen et al., 2004). This resulted in volume maximum projections of ‘microtubules’- and ‘tubulin’-masked densities in cyan, and ‘MIP’-masked densities in yellow. Final visualization and rendering was performed in Blender using 3D scenes imported from UCSF Chimera.

Manual segmentation to obtain protofilament shapes at microtubule ends was performed as described previously (McIntosh et al., 2018; Ogunmolu et al., 2021) using 3dmod (Kremer et al., 1996). Protofilament coordinates were further analyzed using Matlab scripts available at https://github.com/ngudimchuk/Process-PFs.

#### MINFLUX data analysis

Images of microtubule were rendered from MINFLUX data as density map as described in (Schmidt et al., 2021). For the two channel data, the channels were separated by applying cut-off on the “dcr” (detector channel ratio) attribute of the MINFLUX data. The cut-off values were decided by fitting of linear mixture of two Gaussian distributions over the “dcr” values. Data point with “dcr” value in the range between 0 and µ + 0.5σ of the first Gaussian component was considered as belonging to the 1^st^ channel. And data point with “dcr” value in the range between µ - 0.5σ of the second Gaussian component and 1.0 was considered as belonging to the 2^nd^ channel. The rendered MINFLUX data were exported as TIFF images, and used for subsequent analysis in Fiji.

To determine whether the fluorescence signal originated from protein binding on the outside or on the luminal side of the microtubules, we measured the lateral width of the microtubule filaments in the rendered MINFLUX images. To do that, we applied a custom analysis workflow implemented in Fiji. Briefly, for a given microtubule filament, we first extracted the central line along its longitudinal axis. The signal intensity of the nearby regions were then plotted against its distance to the central line, and summed along the length of the filament. Thus we generated the “profile plot”, similar to ImageJ’s intensity profile plot, for each microtubule filament. We then extracted the Full-Width at Half-Maximum (FWHM) of the intensity profile plot, as the estimation of the width of the given filament. A first Fiji script was created to automatically generate the central line segments from the renderend MINFLUX images. In short, it applies line and curvilinear filter to the images to generate first a microtubule segmentation, and then extracts the skeleton of each segmented microtubule filaments as the central line segments. Some manual correction can be applied here to remove the bad segmentation or line segments results, but in most cases it was not necessary. A second Fiji script was then applied to measure the FWHM of the intensity profile of each of the filaments. It reports the FWHM, as well as the length of each filaments, and summarize the results of all filaments, to facilitate further statistical analysis.

### Statistical analysis

All experiments were conducted at least twice. Statistical analysis was performed using GraphPad Prism 9. Statistical details of each experiment, including the statistical tests used, explanation and number of measurement and precision measures can be found in the figure legends.

## Legends to Supplementary Figures and Videos

**Supplementary Figure S1, related to Figure 1.**
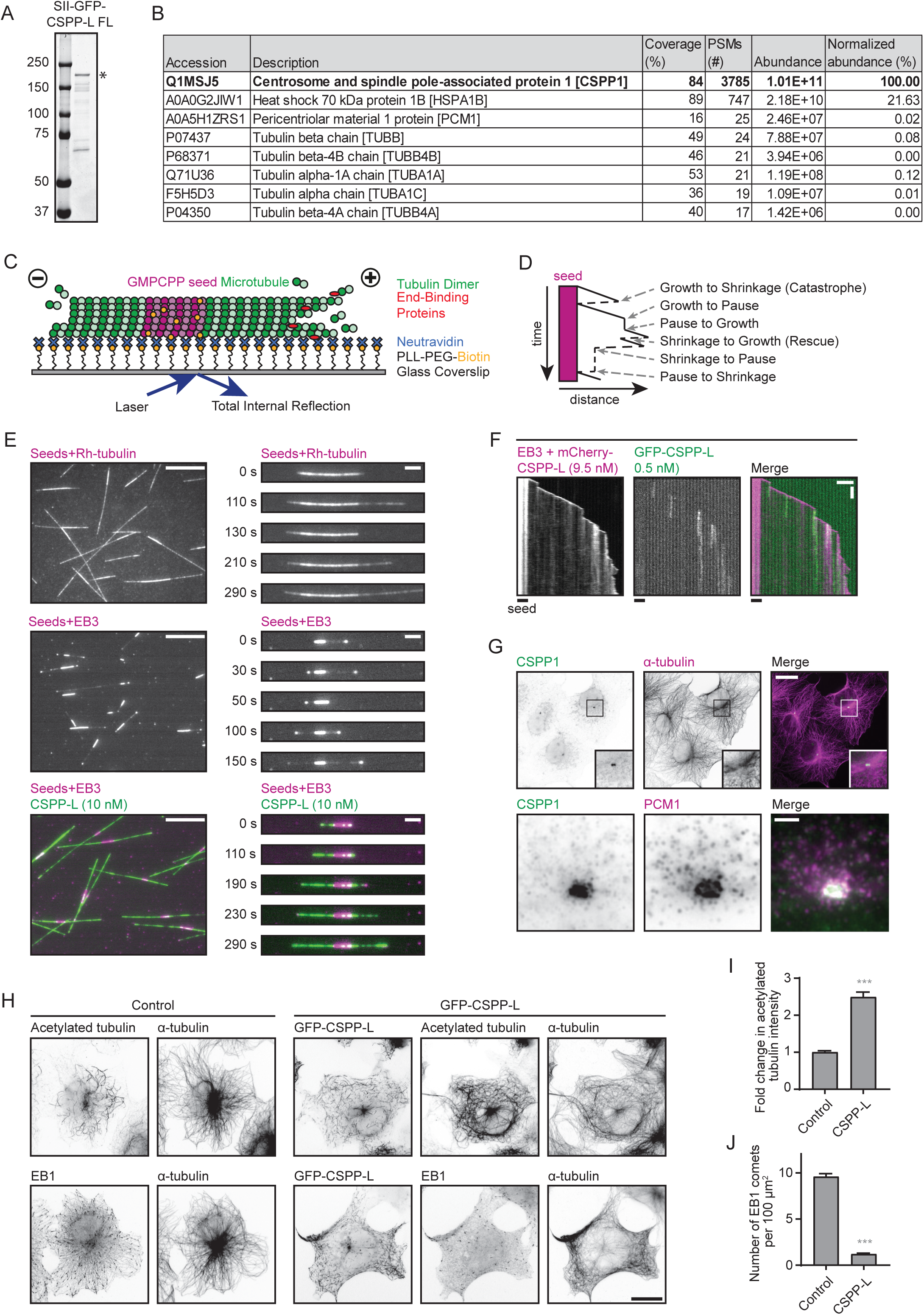
: **Characterization of GFP-CSPP-L in vitro and in COS-7 cells.** A. Analysis of purified GFP-CSPP-L by SDS-PAGE. Asterisk indicates the full-length protein band. Protein concentrations were determined from BSA standard. B. Mass spectrometry analysis of purified GFP-CSPP-L. C. Schematic representation of the in vitro reconstitution assays with dynamic microtubules for imaging with TIRF microscopy. GMPCPP-stabilized microtubule seeds containing fluorescent tubulin, such as rhodamine tubulin (for visualization) and biotinylated tubulin (for surface attachment via NeutrAvidin), are immobilized on a plasma-cleaned coverslip coated with biotinylated poly(L- lysine)-[g]-poly(ethylene glycol) (PLL-PEG-biotin), which is coupled to NeutrAvidin. Microtubule growth from GMPCPP-stabilized seeds is initiated and visualized by the addition of tubulin supplemented with fluorescently-labeled tubulin, or by the addition of unlabeled tubulin combined with fluorescently-tagged EB3. Microtubule plus- and minus-ends are indicated. D. Schematic representation of a kymograph visualizing the various transition events observed and quantified in this paper. E. Field of view (left, scale bar 10 µm) and time-lapse images (right, scale bar 2 µm) illustrating microtubule growth from GMPCPP stabilized microtubule seeds in the presence of 15 µM tubulin supplemented with 3% rhodamine-labelled tubulin, or with 20 nM mCherry-EB3 in presence or absence of 10 nM GFP-CSPP-L. See also Fig. 1B. F. Kymographs illustrating microtubule growth with 20 nM GFP-EB3 together with 9.5 nM mCherry- CSPP-L and 0.5 nM GFP-CSPP-L. Scale bars, 2 μm (horizontal) and 60 s (vertical). G. Widefield fluorescence image of COS-7 cells stained for CSPP1 and α-tubulin or PCM1. Top scale bar, 25 μm; bottom scale bar, 2 µm. H. Widefield fluorescence images of COS-7 cells overexpressing GFP-CSPP-L and stained for α- tubulin and acetylated tubulin or EB1. Scale bar, 20 μm. I. Quantification of mean acetylated tubulin intensity per COS-7 cell (from images as in H). The average mean intensity of cells overexpressing GFP-CSPP-L was normalized to the average mean intensity in control cells. Number of cells analyzed: control cells, n=137; cells overexpressing GFP- CSPP-L, n=77. Data quantified from two experiments. J. Quantification of the number of EB1 comets per 100 µm^2^ in COS-7 cells (from images as in H). Number of cells analyzed: control cells, n=111; cells overexpressing GFP-CSPP-L, n=75. Data quantified from two experiments. For all plots. Error bars represent SEM. Data quantified from two experiments. ***, p< 0.001; Mann- Whitney test.

**Supplementary Figure S2, related to Figure 3:**
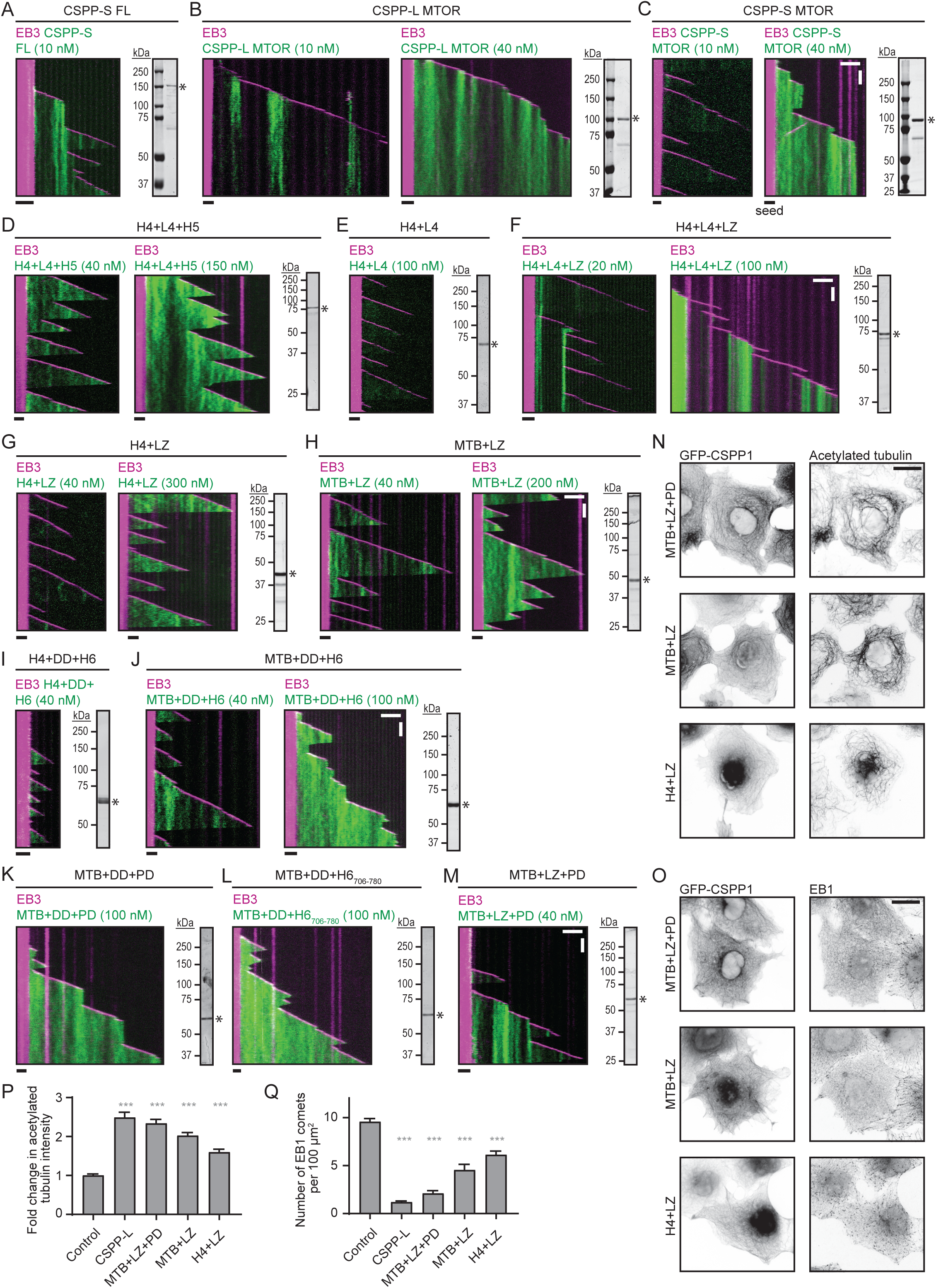
**Shorter CSPP1 constructs are less potent in stabilizing microtubules in cells** A-M. Kymographs of microtubule growth with 20 nM mCherry-EB3 together with the indicated GFP-CSPP1 constructs at the indicated concentrations. Scale bars, 2 μm (horizontal) and 60 s (vertical). Images of SDS-PAGE gels with purified proteins are included for each construct. Asterisk indicates full-length protein band. N-O. Widefield fluorescence images of COS-7 cells overexpressing GFP-CSPP-L and stained for α- tubulin and acetylated tubulin (N) or EB1 (O). Scale bar, 20 μm. P-Q. Quantification of mean acetylated tubulin intensity (P) or quantification of number of EB1 comets per 100 µm^2^ (Q) per COS-7 cell (from images as in N and O). Quantification and statistics as in S1I. Number of cells analyzed acetylated tubulin, EB1: control cells, n=137, n=111; cells overexpressing GFP-CSPP-L, n=77, n=75; cells overexpressing GFP-MTB+LZ+PD, n=83, n=72; cells overexpressing GFP-MTB+LZ, n=70, n=61; cells overexpressing GFP-H4+LZ, n=50, n=75; Data for control and GFP-CSPP-L is the same as in Fig. S1I. Data quantified from two experiments.

**Supplementary Figure S3, related to Figure 4:**
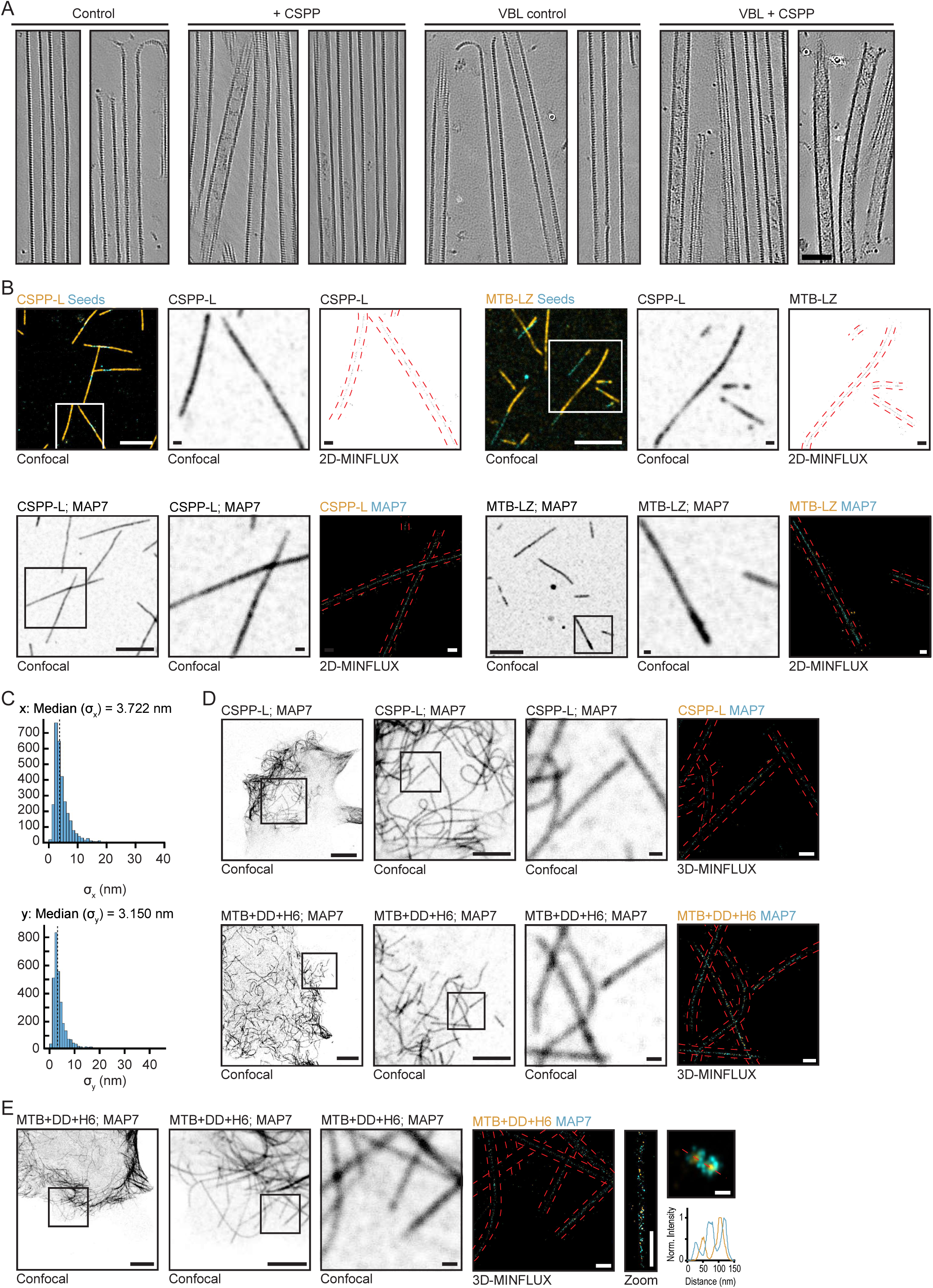
**Characterization of CSPP1 inside the microtubule lumen.** A. Additional tomograms of dynamic microtubules polymerized from GMPCPP-stabilized seeds in the presence or absence of 10 nM GFP-CSPP-L, with or without 250 nM Vinblastine vitrified on EM grids. Scale bar, 50 nm. See also Fig. 4A. B. Additional zooms of confocal images of in vitro MINFLUX regions shown in Fig. 4D-E, G-H. Large confocal field of view (FOV) and 2D-MINFLUX images are identical to the images in Fig. 4D-E, G- H. Panel representation as in (Fig. 4D-E). Scale bars; 5 μm (Large FOV confocal image), 500 nm (Zoom FOV confocal image and 2D-MINFLUX image). C. Standard deviation histograms (x and y-axis) of groups of ≥4 successive localizations from the same fluorophore depicting the localization precision of one example of a single color 2D-MINFLUX measurement of in vitro microtubules polymerized in presence of SNAP-CSPP-L. The localization precision of all single color 2D-MINFLUX measurements used for the FWHM analysis were in the range of 3.0 and 4.3 nm. D. Additional zooms of confocal images of cellular MINFLUX regions shown in Fig. 4I-J. Large confocal field of view (FOV) and 3D-MINFLUX images are identical to the images in Fig. 4I-J. Panel representation as in (Fig. 4I-J). Scale bars; 10 μm (Large FOV confocal image), 5 μm (Medium FOV confocal image), 500 nm (Small FOV confocal image and 3D-MINFLUX image). E. Dual color 3D-MINFLUX measurements of COS-7 cells overexpressing GFP-MAP7 together with SNAP-MTB-DD-H6. Panel representation as in Fig. 4I. Scale bars; 10 μm (Large FOV confocal image), 5 μm (Medium FOV confocal image), 500 nm (Small FOV confocal image, 3D-MINFLUX image and Zoom), 50 nm (maximum intensity projection image).

**Supplementary Figure S4, related to Figure 6.**
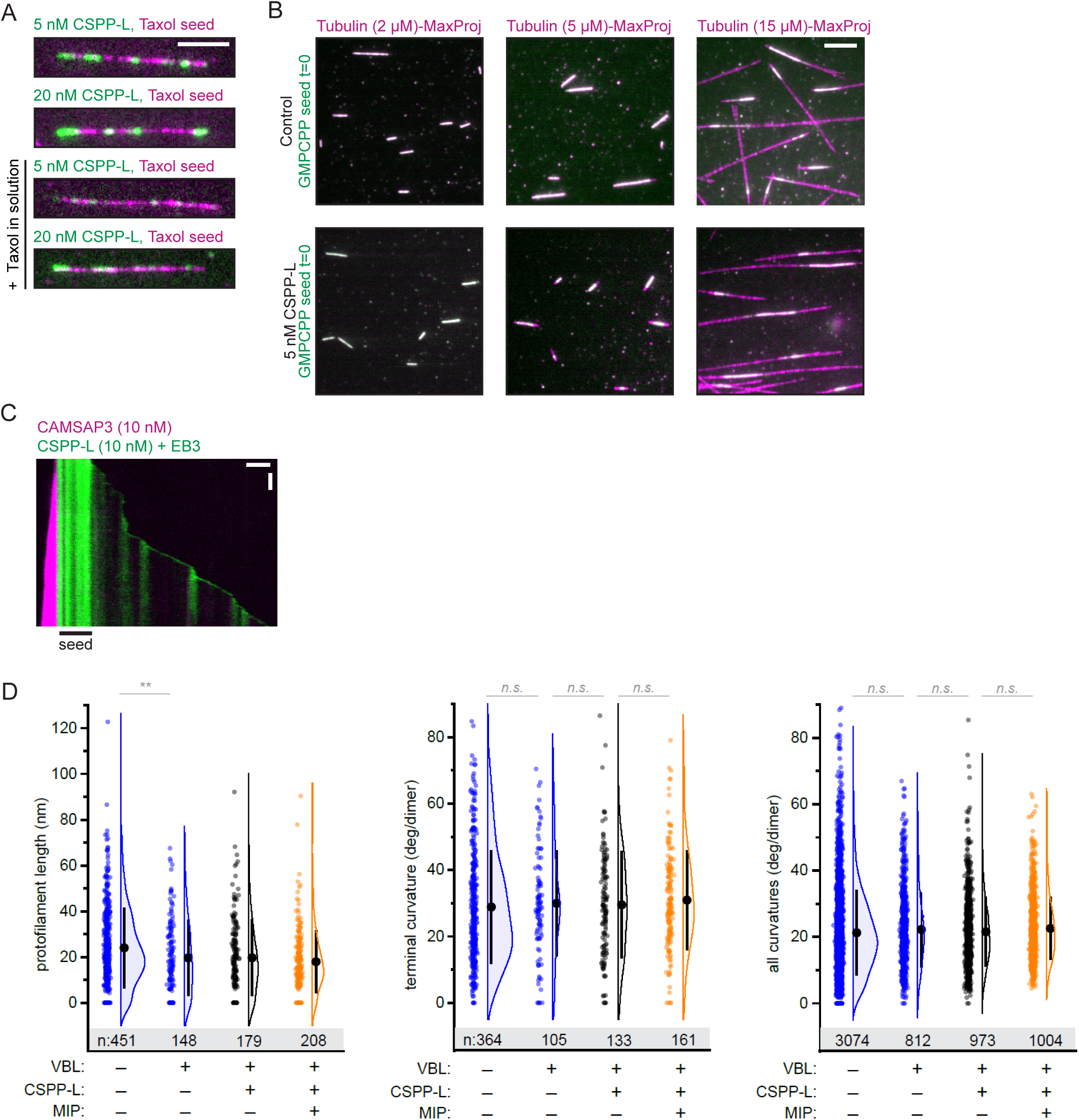
: **CSPP1 stabilized microtubules but there is no change in protofilament length or curvature.** A. Still images from assays with Taxol-stabilized microtubule seeds in the absence or presence 40 µM Taxol and the indicated GFP-CSPP-L concentrations. Scale bar, 5 μm. See also Fig. 6A. B. Microtubule outgrowth from GMPCPP stabilized microtubule seeds five minutes after flowing in 5 nM GFP-CSPP-L at the indicated tubulin concentrations. The first frame of the acquisition (green) is overlayed with the maximum projections of 5 min acquisition (magenta) illustrating the newly grown microtubule lattice. Scale bar 5 µm. C. Kymograph illustrating microtubule growth in the presence of 20 nM mCherry-EB3 together with 10 nM mCherry-CSPP-L and 10 nM GFP-CAMSAP3. Scale bars, 2 μm (horizontal) and 60 s (vertical). D. Quantification of protofilament length, terminal curvature and total curvature (from tomograms as shown in 1K). Blue, orange and grey dots (single data points, tomograms), black circle (mean), SD (error bars). **, p<0.01, n.s., not significant, Mann-Whitney test. Analysis from two experiments.

**Supplementary Video S1. Dynamics of microtubules growing in vitro in the presence CSPP-L.**

An example of TIRF microscopy imaging of microtubules growing from GMPCPP-stabilized microtubule seeds, in presence of 15 µM tubulin (supplemented with 3% rhodamine-tubulin) and 10 nM GFP-CSPP-L.

**Supplementary Video S2. Dynamics of microtubules labeled with CSPP-L and EB3 in COS-7 cells.**

An example of TIRF microscopy imaging of microtubules in COS-7 cells overexpressing GFP- CSPP-L and EB3-mCherry. Full field of view (top) and three enlarged areas of interest are shown (bottom, indicated by white boxes). Arrowheads point to events of interest.

**Supplementary Video S3. 3D view of microtubules in the presence of CSPP-L.**

Rendering of a tomogram acquired with Cryo-ET of microtubules grown in vitro in the presence of GFP-CSPP-L. The denoised densities were segmented into tubulin and microtubules (blue) and all other densities (orange) as described in Methods.

**Supplementary Video S4. 3D view of a microtubule showing MAP7 as an outside ring with CSPP1 construct MTB+DD+H6 inside.**

An example of dual-color 3D-MINFLUX acquisition in COS-7 cells overexpressing SNAP- MTB+DD+H6 and GFP-MAP7.

**Supplementary Video S5. Photodamage of microtubules in COS-7 cells overexpressing GFP- CSPP-L and β-tubulin-mCherry.**

An example of spinning disk confocal imaging of a photodamage experiment in COS-7 cells overexpressing GFP-CSPP-L and β-tubulin-mCherry. Arrowheads point to damage events.

## Key Resources Table

| REAGENT or RESOURCE | SOURCE | IDENTIFIER |
| --- | --- | --- |
| **Antibodies** |  |  |
| Mouse anti-acetylated tubulin | Sigma-Aldrich | Cat# T7451 |
| Purified Mouse Anti-EB1 (Clone 5/EB1) | BD Biosciences | Cat# 610535 |
| Rat anti-tyrosinated α-tubulin (Clone YL1/2) | Abcam | Cat# ab6160 |
| Rabbit anti-CSPP1 | Proteintech | Cat# 11931-1-AP |
| Goat anti-PCM1 antibody (G-6) | Santa Cruz Biotechnology | Cat# sc-398365 |
| Goat anti-Mouse IgG (H+L), Highly Cross-Adsorbed, Alexa Fluor 594 | Thermo Fisher Scientific | Cat# A-11032 |
| Goat anti-Rabbit IgG (H+L), Highly Cross-Adsorbed, Alexa Fluor 488 | Thermo Fisher Scientific | Cat# A-11034 |
| Goat Anti-Rat IgG H&L (Alexa Fluor® 405) preadsorbed | Abcam | Cat# ab175673 |
| Donkey anti-Goat IgG (H+L), Cross-Absorbed, Alexa Fluor 568 | Thermo Fisher Scientific | Cat# A-11057 |
| Donkey anti-Rabbit IgG (H+L), Highly Cross-Absorbed, Alexa Fluor 488 | Thermo Fisher Scientific | Cat# A-21206 |
| Manually conjugated CF680-GFP-Nanobody | Biotium (CF680-dye) + NanoTag (FluoTag®-X4 anti-GFP) | Cat# 92029 + Cat# N0304 |
| FluoTag®-X4 anti-GFP-Alexa 647 | NanoTag | Cat# N0304-AF647-L |
| **Chemicals, Peptides, and Recombinant Proteins** |  |  |
| StrepTactin Sepharose High Performance | GE Healthcare | Cat# 28-9355-99 |
| Polyethyleneimine | Polysciences | Cat# 24765-2 |
| cOmplete™, EDTA-free Protease Inhibitor Cocktail | Roche | Cat# 4693116001 |
| Tubulin protein: porcine brain | Cytoskeleton | Cat# T240 |
| Tubulin protein (fluorescent HiLyte 488): porcine brain | Cytoskeleton | Cat# TL488M |
| Tubulin protein (rhodamine): porcine brain | Cytoskeleton | Cat# TL590M |
| Tubulin protein (biotin labeled): porcine brain | Cytoskeleton | Cat# T333P |
| PLL-PEG-biotin | Susos AG, Switzerland | PLL(20)-g[3.5]-PEG(2)/PEG(3.4)-biotin(50%) |
| Methyl cellulose, 4000 cp | Sigma-Aldrich | Cat# M0512 |
| GMPCPP | Jena Biosciences | Cat# NU-405L |
| GTP | Sigma-Aldrich | Cat# G8877 |
| Glucose oxidase | Sigma-Aldrich | Cat# G7141 |
| Catalase | Sigma-Aldrich | Cat# C9322 |
| DTT | Sigma-Aldrich | Cat# R0861 |
| k-casein | Sigma-Aldrich | Cat# C0406 |
| Neutravidin | Invitrogen | Cat# A-2666 |
| d-Desthiobiotin | Sigma-Aldrich | Cat# D1411 |
| Taxol | Sigma-Aldrich | Cat# T7402 |
| Vinblastine sulfate salt | Sigma-Aldrich | Cat# V1377 |
| Fchitax-3 | (Diaz et al., 2000) | N/A |
| Glucose Oxidase (GLOX-buffer) | Sigma-Aldrich | Cat# G2133 |
| Catalase (GLOX-buffer) | Sigma-Aldrich | Cat# C40 |
| MEA | Sigma-Aldrich | Cat# M9768 |
| StrepII-GFP-CSPP-L full length (CSPP-L FL) | This study | N/A |
| StrepII-GFP-CSPP-L 295-653<sup>^</sup>705-1221 (CSPP-S FL) | This study | N/A |
| StrepII-GFP-CSPP-L 295-849 (CSPP-L MTOR) | This study | N/A |
| StrepII-GFP-CSPP-L 295-653<sup>^</sup>705-849 (CSPP-S MTOR) | This study | N/A |
| StrepII-GFP-CSPP-L 295-653 (H4+L4+H5) | This study | N/A |
| StrepII-GFP-CSPP-L 295-594 (H4+L4) | This study | N/A |
| StrepII-GFP-CSPP-L 295-594+LZ (H4+L4+LZ) | This study | N/A |
| StrepII-GFP-CSPP-L 375-412+(G<sub>4</sub>S)<sub>2</sub>+LZ (H4+LZ) | This study | N/A |
| StrepII-GFP-CSPP-L 375-453+LZ (MTB+LZ) | This study | N/A |
| StrepII-GFP-CSPP-L 375-422<sup>^</sup>583-653<sup>^</sup>705-849 (H4+DD+H6) | This study | N/A |
| StrepII-GFP-CSPP-L 375-453<sup>^</sup>583-653<sup>^</sup>705-849 (MTB+DD+H<sub>6706-780</sub>) | This study | N/A |
| StrepII-GFP-CSPP-L 375-453<sup>^</sup>583-653<sup>^</sup>705-796 (MTB+DD+PD) | This study | N/A |
| StrepII-GFP-CSPP-L 375-453<sup>^</sup>583-653<sup>^</sup>705-780 (MTB+DD+H<sub>6706-780</sub>) | This study | N/A |
| StrepII-GFP-CSPP-L 375-453+LZ+705-796 (MTB+LZ+PD) | This study | N/A |
| StrepII-GFP-MCAK | (Aher et al., 2018) |  |
| mCherry-EB3 full length | (Montenegro Gouveia et al., 2010) | N/A |
| GFP-EB3 full length | (Montenegro Gouveia et al., 2010) | N/A |
| StrepII-GFP-MAP7 N-terminus | (Hooikaas et al., 2019) | N/A |
| FuGENE6 | Promega | Cat# E2691 |
| SNAP-Surface® Alexa Fluor® 647 | NEB | Cat# S9136S |
| SNAP-Abberior FLUX 680 | Abberior | FX680 |
| Glutaraldehyde | Electron Microscopy Sciences | Cat# #16110 |
| Lacey EM grids | SPI supplies | Cat# 3840G |
| 5 nm gold particles | Sigma-Aldrich | Cat# 808628 |
| Spherical Gold Nanoparticles (200 nm) | Nanopartz | Cat# A11-200-CIT-DIH-1-10 |
| Experimental Models: Cell Lines |  |  |
| Human: HEK293T | ATCC | CRL-11268 |
| Monkey: COS-7 | ATCC | CRL-1651 |
| Recombinant DNA |  |  |
| Human EB3-mCherry | (Stepanova et al., 2003) | N/A |
| Human βIVb-tubulin-mCherry | (Bouchet et al., 2016) | N/A |
| Human StrepII-GFP-MAP7 | (Hooikaas et al., 2019) | N/A |
| Software and Algorithms |  |  |
| ImageJ | NIH | https://imagej.nih.gov/ij/ |
| Metamorph Version 7.8 | Molecular Devices | https://www.moleculardevices.com/products/cellular-imaging-systems/acquisition-and-analysis-software/metamorph-microscopy |
| GraphPad Prism 9 | GraphPad Software | https://www.graphpad.com/scientific-software/prism/ |
| KymoResliceWide plugin | Eugene Katrukha | https://github.com/ekatrunkha/KymoResliceWide |
| DoM Utrecht plugin | Eugene Katrukha | https://github.com/ekatrunkha/DoM_Utrecht |
| MATLAB | MathWorks | https://www.mathworks.com/ |
| MATLAB code for max intensity fit of EB3 comet and AUC of fitted CSPP-L signal | (Rai et al., 2020) | N/A |
| IMOD v. 4.11 | (Kremer et al., 1996) | https://bio3d.colorado.edu/imod/ |
| EMAN 2, tomoseg | (Chen et al., 2017) | https://cryoem.bcm.edu/cryoem/downloads/view_eman2_versions |
| Python scripts for cryoCARE denoising | (Buchholz et al., 2019; Ogunmolu et al., 2021) | https://github.com/NemoAndrea/cryoCARE-hpc04 |
| MATLAB scripts for analysis of protofilament shapes | This study | https://github.com/ngudimchuk/Process-PFs |
| UCSF Chimera | (Pettersen et al., 2004) | https://www.cgl.ucsf.edu/chimera/download.html |
| MATLAB and Fiji Groovy scripts for MINFLUX Analysis | This study | https://github.com/EMBL-ICLM/microtubule_width_measurement_MINFLUX |

## References

Aher, A., Kok, M., Sharma, A., Rai, A., Olieric, N., Rodriguez-Garcia, R., Katrukha, E.A., Weinert, T., Olieric, V., Kapitein, L.C., et al. (2018). CLASP Suppresses Microtubule Catastrophes through a Single TOG Domain. Dev Cell 46, 40–58.

Aher, A., Rai, D., Schaedel, L., Gaillard, J., John, K., Liu, Q., Altelaar, M., Blanchoin, L., Thery, M., and Akhmanova, A. (2020). CLASP Mediates Microtubule Repair by Restricting Lattice Damage and Regulating Tubulin Incorporation. Curr Biol 30, 2175–2183.

Akhmanova, A., and Steinmetz, M.O. (2015). Control of microtubule organization and dynamics: two ends in the limelight. Nat Rev Mol Cell Biol 16, 711–726.

Akizu, N., Silhavy, J.L., Rosti, R.O., Scott, E., Fenstermaker, A.G., Schroth, J., Zaki, M.S., Sanchez, H., Gupta, N., Kabra, M., et al. (2014). Mutations in CSPP1 lead to classical Joubert syndrome. Am J Hum Genet 94, 80–86.

Al-Jassar, C., Andreeva, A., Barnabas, D.D., McLaughlin, S.H., Johnson, C.M., Yu, M., and van Breugel, M. (2017). The Ciliopathy-Associated Cep104 Protein Interacts with Tubulin and Nek1 Kinase. Structure 25, 146–156.

Andreu-Cervera, A., Catala, M., and Schneider-Maunoury, S. (2021). Cilia, ciliopathies and hedgehog-related forebrain developmental disorders. Neurobiol Dis 150, 105236.

Arnal, I., and Wade, R.H. (1995). How does taxol stabilize microtubules? Curr Biol 5, 900–908.

Asiedu, M., Wu, D., Matsumura, F., and Wei, Q. (2009). Centrosome/spindle pole-associated protein regulates cytokinesis via promoting the recruitment of MyoGEF to the central spindle. Mol Biol Cell 20, 1428–1440.

Aumeier, C., Schaedel, L., Gaillard, J., John, K., Blanchoin, L., and Thery, M. (2016). Self-repair promotes microtubule rescue. Nat Cell Biol 18, 1054–1064.

Balzarotti, F., Eilers, Y., Gwosch, K.C., Gynna, A.H., Westphal, V., Stefani, F.D., Elf, J., and Hell, S.W. (2017). Nanometer resolution imaging and tracking of fluorescent molecules with minimal photon fluxes. Science 355, 606–612.

Basnet, N., Nedozralova, H., Crevenna, A.H., Bodakuntla, S., Schlichthaerle, T., Taschner, M., Cardone, G., Janke, C., Jungmann, R., Magiera, M.M., et al. (2018). Direct induction of microtubule branching by microtubule nucleation factor SSNA1. Nat Cell Biol 20, 1172–1180.

Bieling, P., Laan, L., Schek, H., Munteanu, E.L., Sandblad, L., Dogterom, M., Brunner, D., and Surrey, T. (2007). Reconstitution of a microtubule plus-end tracking system in vitro. Nature 450, 1100–1105.

Bouchet, B.P., Noordstra, I., van Amersfoort, M., Katrukha, E.A., Ammon, Y.C., Ter Hoeve, N.D., Hodgson, L., Dogterom, M., Derksen, P.W., and Akhmanova, A. (2016). Mesenchymal Cell Invasion Requires Cooperative Regulation of Persistent Microtubule Growth by SLAIN2 and CLASP1. Dev Cell 39, 708–723.

Buchholz, T.O., Krull, A., Shahidi, R., Pigino, G., Jekely, G., and Jug, F. (2019). Content-aware image restoration for electron microscopy. Methods Cell Biol 152, 277–289.

Chen, M., Dai, W., Sun, S.Y., Jonasch, D., He, C.Y., Schmid, M.F., Chiu, W., and Ludtke, S.J. (2017). Convolutional neural networks for automated annotation of cellular cryo-electron tomograms. Nat Methods 14, 983–985.

Cuveillier, C., Delaroche, J., Seggio, M., Gory-Faure, S., Bosc, C., Denarier, E., Bacia, M., Schoehn, G., Mohrbach, H., Kulic, I., et al. (2020). MAP6 is an intraluminal protein that induces neuronal microtubules to coil. Sci Adv 6, eaaz4344.

Das, A., Dickinson, D.J., Wood, C.C., Goldstein, B., and Slep, K.C. (2015). Crescerin uses a TOG domain array to regulate microtubules in the primary cilium. Mol Biol Cell 26, 4248–4264.

Desai, A., and Mitchison, T.J. (1997). Microtubule polymerization dynamics. Annu Rev Cell Dev Biol 13, 83–117.

Diaz, J.F., Strobe, R., Engelborghs, Y., Souto, A.A., and Andreu, J.M. (2000). Molecular recognition of taxol by microtubules. Kinetics and thermodynamics of binding of fluorescent taxol derivatives to an exposed site. J Biol Chem 275, 26265–26276.

Elie-Caille, C., Severin, F., Helenius, J., Howard, J., Muller, D.J., and Hyman, A.A. (2007). Straight GDP-tubulin protofilaments form in the presence of taxol. Curr Biol 17, 1765–1770.

Ferro, L.S., Fang, Q., Eshun-Wilson, L., Fernandes, J., Jack, A., Farrell, D.P., Golcuk, M., Huijben, T., Costa, K., Gur, M., et al. (2022). Structural and functional insight into regulation of kinesin-1 by microtubule-associated protein MAP7. Science 375, 326–331.

Frikstad, K.M., Molinari, E., Thoresen, M., Ramsbottom, S.A., Hughes, F., Letteboer, S.J.F., Gilani, S., Schink, K.O., Stokke, T., Geimer, S., et al. (2019). A CEP104-CSPP1 Complex Is Required for Formation of Primary Cilia Competent in Hedgehog Signaling. Cell Rep 28, 1907–1922.

Gazzola, M., Schaeffer, A., Vianay, B., Gaillard, J., Blanchoin, L., and Théry, M. (2022). Microtubules self-repair in living cells. bioRxiv, 2022.2003.2031.486545.

Gell, C., Bormuth, V., Brouhard, G.J., Cohen, D.N., Diez, S., Friel, C.T., Helenius, J., Nitzsche, B., Petzold, H., Ribbe, J., et al. (2010). Microtubule dynamics reconstituted in vitro and imaged by single-molecule fluorescence microscopy. Methods Cell Biol 95, 221–245.

Gudimchuk, N.B., and McIntosh, J.R. (2021). Regulation of microtubule dynamics, mechanics and function through the growing tip. Nat Rev Mol Cell Biol 22, 777–795.

Gui, M., Farley, H., Anujan, P., Anderson, J.R., Maxwell, D.W., Whitchurch, J.B., Botsch, J.J., Qiu, T., Meleppattu, S., Singh, S.K., et al. (2021). De novo identification of mammalian ciliary motility proteins using cryo-EM. Cell 184, 5791–5806.

Gwosch, K.C., Pape, J.K., Balzarotti, F., Hoess, P., Ellenberg, J., Ries, J., and Hell, S.W. (2020). MINFLUX nanoscopy delivers 3D multicolor nanometer resolution in cells. Nat Methods 17, 217–224.

Hildebrandt, F., Benzing, T., and Katsanis, N. (2011). Ciliopathies. N Engl J Med 364, 1533–1543.

Hooikaas, P.J., Martin, M., Muhlethaler, T., Kuijntjes, G.J., Peeters, C.A.E., Katrukha, E.A., Ferrari, L., Stucchi, R., Verhagen, D.G.F., van Riel, W.E., et al. (2019). MAP7 family proteins regulate kinesin-1 recruitment and activation. J Cell Biol 218, 1298–1318.

Ichikawa, M., and Bui, K.H. (2018). Microtubule Inner Proteins: A Meshwork of Luminal Proteins Stabilizing the Doublet Microtubule. Bioessays 40, 1700209.

Jiang, K., Toedt, G., Montenegro Gouveia, S., Davey, N.E., Hua, S., van der Vaart, B., Grigoriev, I., Larsen, J., Pedersen, L.B., Bezstarosti, K., et al. (2012). A Proteome-wide screen for mammalian SxIP motif-containing microtubule plus-end tracking proteins. Curr Biol 22, 1800–1807.

Jumper, J., Evans, R., Pritzel, A., Green, T., Figurnov, M., Ronneberger, O., Tunyasuvunakool, K., Bates, R., Zidek, A., Potapenko, A., et al. (2021). Highly accurate protein structure prediction with AlphaFold. Nature 596, 583–589.

Komarova, Y., De Groot, C.O., Grigoriev, I., Gouveia, S.M., Munteanu, E.L., Schober, J.M., Honnappa, S., Buey, R.M., Hoogenraad, C.C., Dogterom, M., et al. (2009). Mammalian end binding proteins control persistent microtubule growth. J Cell Biol 184, 691–706.

Kremer, J.R., Mastronarde, D.N., and McIntosh, J.R. (1996). Computer visualization of three-dimensional image data using IMOD. J Struct Biol 116, 71–76.

Latour, B.L., Van De Weghe, J.C., Rusterholz, T.D., Letteboer, S.J., Gomez, A., Shaheen, R., Gesemann, M., Karamzade, A., Asadollahi, M., Barroso-Gil, M., et al. (2020). Dysfunction of the ciliary ARMC9/TOGARAM1 protein module causes Joubert syndrome. J Clin Invest 130, 4423–4439.

Lawrence, E.J., Arpag, G., Arnaiz, C., and Zanic, M. (2021). SSNA1 stabilizes dynamic microtubules and detects microtubule damage. Elife 10, e67282.

Ma, M., Stoyanova, M., Rademacher, G., Dutcher, S.K., Brown, A., and Zhang, R. (2019). Structure of the Decorated Ciliary Doublet Microtubule. Cell 179, 909–922.

Magiera, M.M., Singh, P., Gadadhar, S., and Janke, C. (2018). Tubulin Posttranslational Modifications and Emerging Links to Human Disease. Cell 173, 1323–1327.

Mastronarde, D.N. (2005). Automated electron microscope tomography using robust prediction of specimen movements. J Struct Biol 152, 36–51.

McIntosh, J.R., O’Toole, E., Morgan, G., Austin, J., Ulyanov, E., Ataullakhanov, F., and Gudimchuk, N. (2018). Microtubules grow by the addition of bent guanosine triphosphate tubulin to the tips of curved protofilaments. J Cell Biol 217, 2691–2708.

Mimori-Kiyosue, Y., Shiina, N., and Tsukita, S. (2000). The dynamic behavior of the APC-binding protein EB1 on the distal ends of microtubules. Curr Biol 10, 865–868.

Mohan, R., Katrukha, E.A., Doodhi, H., Smal, I., Meijering, E., Kapitein, L.C., Steinmetz, M.O., and Akhmanova, A. (2013). End-binding proteins sensitize microtubules to the action of microtubule-targeting agents. Proc Natl Acad Sci U S A 110, 8900–8905.

Montenegro Gouveia, S., Leslie, K., Kapitein, L.C., Buey, R.M., Grigoriev, I., Wagenbach, M., Smal, I., Meijering, E., Hoogenraad, C.C., Wordeman, L., et al. (2010). In vitro reconstitution of the functional interplay between MCAK and EB3 at microtubule plus ends. Curr Biol 20, 1717–1722.

Odde, D. (1998). Diffusion inside microtubules. Eur Biophys J 27, 514–520.

Ogunmolu, F.E., Moradi, S., Volkov, V.A., van Hoorn, C., Wu, J., Andrea, N., Hua, S., Jiang, K., Vakonakis, I., Potočnjak, M., et al. (2021). Microtubule plus-end regulation by centriolar cap proteins. bioRxiv, 2021.2012.2029.474442.

Patzke, S., Hauge, H., Sioud, M., Finne, E.F., Sivertsen, E.A., Delabie, J., Stokke, T., and Aasheim, H.C. (2005). Identification of a novel centrosome/microtubule-associated coiled-coil protein involved in cell-cycle progression and spindle organization. Oncogene 24, 1159–1173.

Patzke, S., Redick, S., Warsame, A., Murga-Zamalloa, C.A., Khanna, H., Doxsey, S., and Stokke, T. (2010). CSPP is a ciliary protein interacting with Nephrocystin 8 and required for cilia formation. Mol Biol Cell 21, 2555–2567.

Patzke, S., Stokke, T., and Aasheim, H.C. (2006). CSPP and CSPP-L associate with centrosomes and microtubules and differently affect microtubule organization. J Cell Physiol 209, 199–210.

Pettersen, E.F., Goddard, T.D., Huang, C.C., Couch, G.S., Greenblatt, D.M., Meng, E.C., and Ferrin, T.E. (2004). UCSF Chimera--a visualization system for exploratory research and analysis. J Comput Chem 25, 1605–1612.

Prota, A.E., Bargsten, K., Zurwerra, D., Field, J.J., Diaz, J.F., Altmann, K.H., and Steinmetz, M.O. (2013). Molecular mechanism of action of microtubule-stabilizing anticancer agents. Science 339, 587–590.

Rai, A., Liu, T., Glauser, S., Katrukha, E.A., Estevez-Gallego, J., Rodriguez-Garcia, R., Fang, W.S., Diaz, J.F., Steinmetz, M.O., Altmann, K.H., et al. (2020). Taxanes convert regions of perturbed microtubule growth into rescue sites. Nat Mater 19, 355–365.

Rai, A., Liu, T., Katrukha, E.A., Estevez-Gallego, J., Manka, S.W., Paterson, I., Diaz, J.F., Kapitein, L.C., Moores, C.A., and Akhmanova, A. (2021). Lattice defects induced by microtubule-stabilizing agents exert a long-range effect on microtubule growth by promoting catastrophes. Proc Natl Acad Sci U S A 118, e2112261118.

Reiter, J.F., and Leroux, M.R. (2017). Genes and molecular pathways underpinning ciliopathies. Nat Rev Mol Cell Biol 18, 533–547.

Rezabkova, L., Kraatz, S.H., Akhmanova, A., Steinmetz, M.O., and Kammerer, R.A. (2016). Biophysical and Structural Characterization of the Centriolar Protein Cep104 Interaction Network. J Biol Chem 291, 18496–18504.

Schmidt, R., Weihs, T., Wurm, C.A., Jansen, I., Rehman, J., Sahl, S.J., and Hell, S.W. (2021). MINFLUX nanometer-scale 3D imaging and microsecond-range tracking on a common fluorescence microscope. Nat Commun 12, 1478.

Shaheen, R., Shamseldin, H.E., Loucks, C.M., Seidahmed, M.Z., Ansari, S., Ibrahim Khalil, M., Al- Yacoub, N., Davis, E.E., Mola, N.A., Szymanska, K., et al. (2014). Mutations in CSPP1, encoding a core centrosomal protein, cause a range of ciliopathy phenotypes in humans. Am J Hum Genet 94, 73–79.

Sharma, A., Aher, A., Dynes, N.J., Frey, D., Katrukha, E.A., Jaussi, R., Grigoriev, I., Croisier, M., Kammerer, R.A., Akhmanova, A., et al. (2016). Centriolar CPAP/SAS-4 Imparts Slow Processive Microtubule Growth. Dev Cell 37, 362–376.

Steinmetz, M.O., and Prota, A.E. (2018). Microtubule-Targeting Agents: Strategies To Hijack the Cytoskeleton. Trends Cell Biol 28, 776–792.

Stepanova, T., Slemmer, J., Hoogenraad, C.C., Lansbergen, G., Dortland, B., De Zeeuw, C.I., Grosveld, F., van Cappellen, G., Akhmanova, A., and Galjart, N. (2003). Visualization of microtubule growth in cultured neurons via the use of EB3-GFP (end-binding protein 3-green fluorescent protein). J Neurosci 23, 2655–2664.

Triclin, S., Inoue, D., Gaillard, J., Htet, Z.M., DeSantis, M.E., Portran, D., Derivery, E., Aumeier, C., Schaedel, L., John, K., et al. (2021). Self-repair protects microtubules from destruction by molecular motors. Nat Mater 20, 883–891.

Tuz, K., Bachmann-Gagescu, R., O’Day, D.R., Hua, K., Isabella, C.R., Phelps, I.G., Stolarski, A.E., O’Roak, B.J., Dempsey, J.C., Lourenco, C., et al. (2014). Mutations in CSPP1 cause primary cilia abnormalities and Joubert syndrome with or without Jeune asphyxiating thoracic dystrophy. Am J Hum Genet 94, 62–72.

van Riel, W.E., Rai, A., Bianchi, S., Katrukha, E.A., Liu, Q., Heck, A.J., Hoogenraad, C.C., Steinmetz, M.O., Kapitein, L.C., and Akhmanova, A. (2017). Kinesin-4 KIF21B is a potent microtubule pausing factor. Elife 6, e24746.

Varadi, M., Anyango, S., Deshpande, M., Nair, S., Natassia, C., Yordanova, G., Yuan, D., Stroe, O., Wood, G., Laydon, A., et al. (2022). AlphaFold Protein Structure Database: massively expanding the structural coverage of protein-sequence space with high-accuracy models. Nucleic Acids Res 50, D439–D444.

Vemu, A., Szczesna, E., Zehr, E.A., Spector, J.O., Grigorieff, N., Deaconescu, A.M., and Roll-Mecak, A. (2018). Severing enzymes amplify microtubule arrays through lattice GTP-tubulin incorporation. Science 361, eaau1504.

Wieczorek, M., Bechstedt, S., Chaaban, S., and Brouhard, G.J. (2015). Microtubule-associated proteins control the kinetics of microtubule nucleation. Nat Cell Biol 17, 907–916.

Zheng, S.Q., Palovcak, E., Armache, J.P., Verba, K.A., Cheng, Y., and Agard, D.A. (2017). MotionCor2: anisotropic correction of beam-induced motion for improved cryo-electron microscopy. Nat Methods 14, 331–332.

